# cazy_webscraper: local compilation and interrogation of comprehensive CAZyme datasets

**DOI:** 10.1101/2022.12.02.518825

**Authors:** Emma E. M. Hobbs, Tracey M. Gloster, Leighton Pritchard

**Affiliations:** School of Biology and Biomedical Sciences Research Complex, University of St Andrews, North Haugh, St Andrews, Fife, KY16 9ST, UK; Strathclyde Institute of Pharmacy and Biomedical Sciences, University of Strathclyde, Glasgow, G4 0RE, UK; Cell and Molecular Sciences, James Hutton Institute, Invergowrie, Dundee, DD2 5DA, UK

**Keywords:** Carbohydrate-active enzymes, CAZy, Software, Database

## Abstract

Carbohydrate Active enZymes (CAZymes) are pivotal in biological processes including energy metabolism, cell structure maintenance, signalling and pathogen recognition. Bioinformatic prediction and mining of CAZymes improves our understanding of these activities, and enables discovery of candidates of interest for industrial biotechnology, particularly the processing of organic waste for biofuel production. CAZy (www.cazy.org) is a high-quality, manually-curated and authoritative database of CAZymes that is often the starting point for these analyses. Automated querying, and integration of CAZy data with other public datasets would constitute a powerful resource for mining and exploring CAZyme diversity. However, CAZy does not itself provide methods to automate queries, or integrate annotation data from other sources (except by following hyperlinks) to support further analysis.

To overcome these limitations we developed `cazy_webscraper`, a command-line tool that retrieves data from CAZy and other online resources to build a local, shareable, and reproducible database that augments and extends the authoritative CAZy database. `cazy_webscraper`’s integration of curated CAZyme annotations with their corresponding protein sequences, up to date taxonomy assignments, and protein structure data facilitates automated large-scale and targeted bioinformatic CAZyme family analysis and candidate screening. This tool has found widespread uptake in the community, with over 20,000 downloads.

We demonstrate the use and application of `cazy_webscraper` to: (i) augment, update and correct CAZy database accessions; (ii) explore taxonomic distribution of CAZymes recorded in CAZy, identifying underrepresented taxa and unusual CAZy class distributions; and (iii) investigate three CAZymes having potential biotechnological application for degradation of biomass, but lacking a representative structure in the PDB database. We describe in general how `cazy_webscraper` facilitates functional, structural and evolutionary studies to aid identification of candidate enzymes for further characterisation, and specifically note that CAZy provides supporting evidence for recent expansion of the Auxiliary Activities (AA) CAZy family in eukaryotes, consistent with functions potentially specific to eukaryotic lifestyles.

**Supplementary information:** `cazy_webscraper` source code is available at https://github.com/HobnobMancer/cazy_webscraper, and online documentation is provided at https://cazywebscraper.readthedocs.io.

## Background

Carbohydrate Active enZymes (CAZymes) catalyse modification, synthesis and degradation of polysaccharides and glycoconjugates (1). They are vital in many biological pathways including metabolism and cell signalling, are used for industrial bioprocessing of organic materials, and are of fundamental research interest (2–5).

The most comprehensive and authoritative CAZyme resource is the manually-curated CAZy database (www.cazy.org). CAZy is built by classification of sequenced CAZymes (glycoside hydrolases, GHs; glycosyltransferases, GTs; polysaccharide lyases, PLs; carbohydrate esterases, CEs; auxiliary activities, AAs; and non-catalytic carbohydrate binding modules, CBMs) into sequence similarity-based families corresponding to presumed shared mechanism and structural fold, enabling principled annotation and prediction of enzyme function (6). This database is widely used as the primary reference database for bioinformatic analyses of CAZymes. For many downstream analyses and applications using CAZy data, including genome annotation and the training of machine learning models, it is necessary to be able to filter sequences, often on the basis of additional data held in other resources. However, CAZy currently provides the majority of its data only in two forms: a plain text download containing all members of a single CAZy family; and *via* a browser interface, paginated in small groups for a subset of functionally and/or structurally characterised members of the family. The absence of a public application programming interface (API) or query function makes automated retrieval and querying of large and/or user-filtered datasets from CAZy for bioinformatic analysis more challenging.

Several software tools have been developed to overcome the limitations of the CAZy web interface, but many are no longer supported or have not been updated to reflect recent changes to the CAZy site structure (6). One such deprecated tool is `CAZy_utils` (https://github.com/nielshanson/CAZy_utils) which was similar in function to `cazy_webscraper` in that it also built a comprehensive local CAZyme database from the available CAZy data. Other tools retrieve only limited data from CAZy: `cazyseqs` (https://github.com/walshaw/cazyseqs) gathers only protein sequences from CAZy families, not CAZy family annotations; and `cazy_parser` (7) compiles HTML tables from the CAZy website into a single comma-separated variable (CSV) file.

To fill this gap in capability, and to facilitate downstream analyses using CAZy data, we present `cazy_webscraper` (https://doi.org/10.5281/zenodo.6343936), a command-line tool with significant community uptake (averaging 1,800 downloads per month, February-July 2022 (8)) that automates data retrieval from the CAZy database and other resources, integrating these into a local SQLite3 database. `cazy_webscraper` can integrate protein sequence, taxonomic lineage, Enzyme Commission (EC) annotation and structural information data from public repositories including the National Center for Biotechnology Information (NCBI) (9), UniProt (10), Research Collaboratory for Structural Protein Data Bank (RSCB PDB) (11) and the Genome Taxonomy Database (GTDB) (12). All data retrieval is logged for audit and reproducibility. The database constructed by `cazy_webscraper` is reusable and shareable, enabling replication of downstream bioinformatic analyses. `cazy_webscraper` also provides an API to this database, enabling integration as part of automated analyses.

We demonstrate the use and application of `cazy_webscraper` with practical examples. We survey taxonomic diversity in CAZy, highlighting uneven representation of archaeal lineages and CAZy families to identify scope for strategic investigations to extend our understanding. We analyse sequence diversity across the PL20 family, identifying a conserved enzyme represented only in *Streptomyces*, and a *β*-1,4-glucuronan lyase (EC 4.2.2.14) from *Galbibacter* sp. BG1 that may be the only membrane-associated PL20 glucuronan lyase in CAZy. In addition, we survey the known structural representation of carbohydrate esterases (CEs) in CAZy, and identify and investigate two CAZyme groups of potential interest to industrial biotechnology that currently have no structural representatives in the RCSB PDB.

## Implementation

`cazy_webscraper` is implemented as a Python package that automates retrieval of records from CAZy downloaded as a plain text file, and builds a local SQLite3 database (13) from this (schema shown in figure 1). The object relational mapping (ORM) is specified and managed using SQLAlchemy (14).

**Fig. 1.**
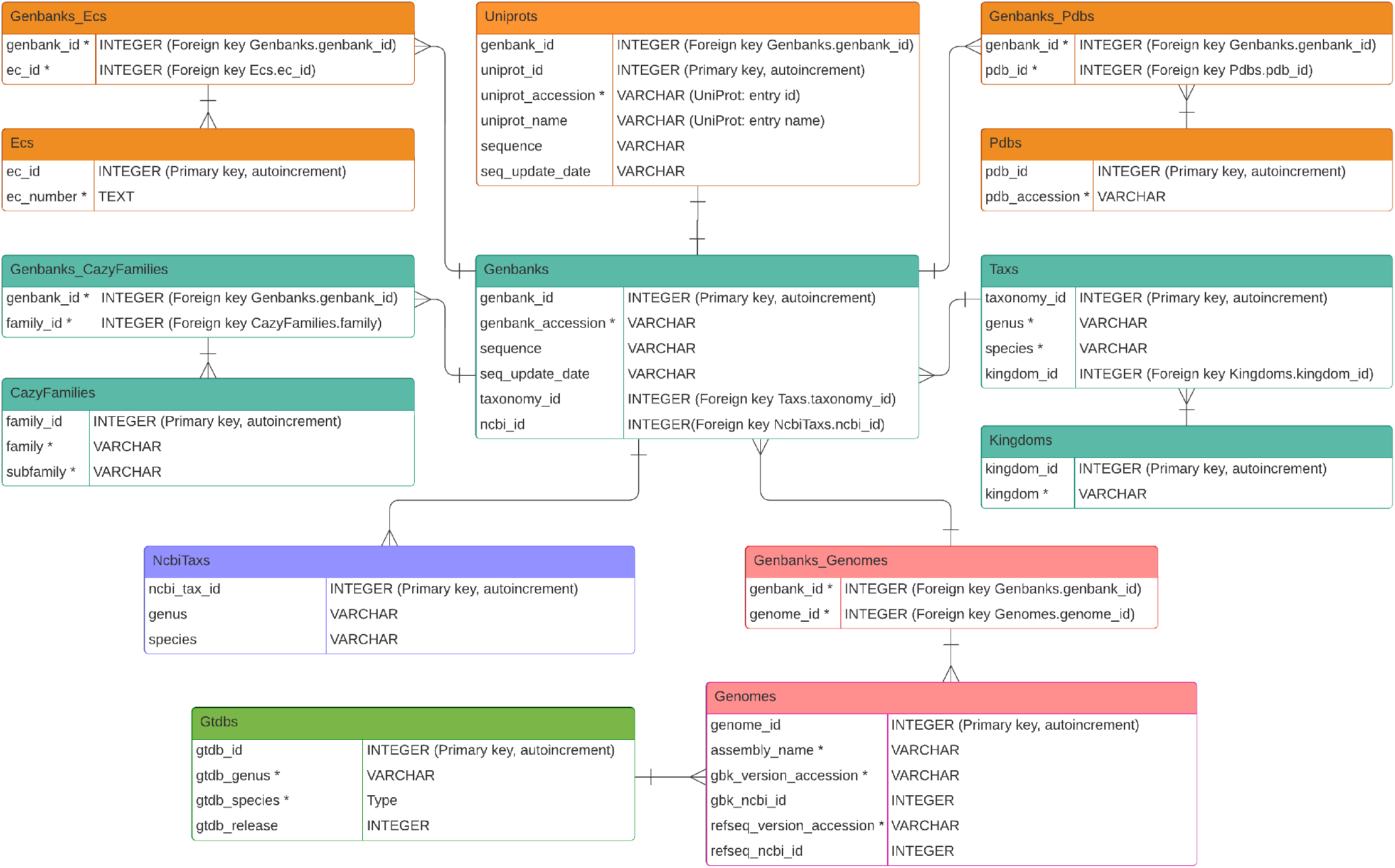
Entity-relationship model of the `cazy_webscraper` database structure. Teal boxes represent tables containing data collected from CAZy. Orange tables contain data retrieved from UniProtKB. Purple and pink boxes represent tables containing data retrieved from NCBI, and the green table contains data retrieved from GTDB. The datatype VARCHAR represents a variable length character string. Fields under UNIQUE constraint are marked with *, and such fields within the same table are under joint UNIQUE constraint.

CAZy provides a plain text file dump of the complete database for public use. This is downloaded by `cazy_webscraper` and cached locally to enable multiple distinct filtering operations to be applied on the original dataset. This facilitates reproduction, and allows for generation of specialised databases, such as a separate database for each CAZy class. The CAZy download can be parsed to extract all CAZyme records, or only records matching user-specified criteria. `cazy_webscraper` imports CAZy fields including GenBank accessions, source organism taxon (genus, species, and kingdom), and CAZy family. At the time of writing, PubMed IDs and clan data (an additional CAZy classification encompassing groups of families that share a fold and catalytic machinery) (6) are not imported as these are not provided in the downloadable database dump.

`cazy_webscraper` identifies distinct CAZymes uniquely by their NCBI accession as recorded in CAZy. These are mostly GenBank accessions, although some entries in CAZy are recorded as RefSeq accessions. As the same CAZyme may be associated with more than one CAZy family or subfamily classification, our approach eliminates some redundancy present in the CAZy dataset.

`cazy_webscraper` will by default construct a new database, but can also update an existing local database. Updating a local database with different filters, or a series of successive data retrieval operations, does not introduce duplicate CAZyme records. `cazy_webscraper` caches unprocessed data as it is retrieved from CAZy, or from external databases such as UniProt, NCBI and GTDB. If a download is interrupted, `cazy_webscraper` can be resumed from the point it halted.

### Filtering CAZy data on import

The publicly-available CAZy database download contains the complete set of CAZyme records, but not all data available at the CAZy web service. `cazy_webscraper` filtering options can be used to import into the local database only records matching combinations of user-specified criteria, including: CAZy class, CAZy family, CAZy subfamily, kingdom, genus, species and/or strain. Filters can be specified using a configuration file or by passing parameters at the command-line. Similar filters, and an additional EC number filter, can be used to control the retrieval of data from external databases such as UniProt, NCBI, PDB and GTDB. Table 1 lists example commands and a summary of the data imported.

**Table 1.**
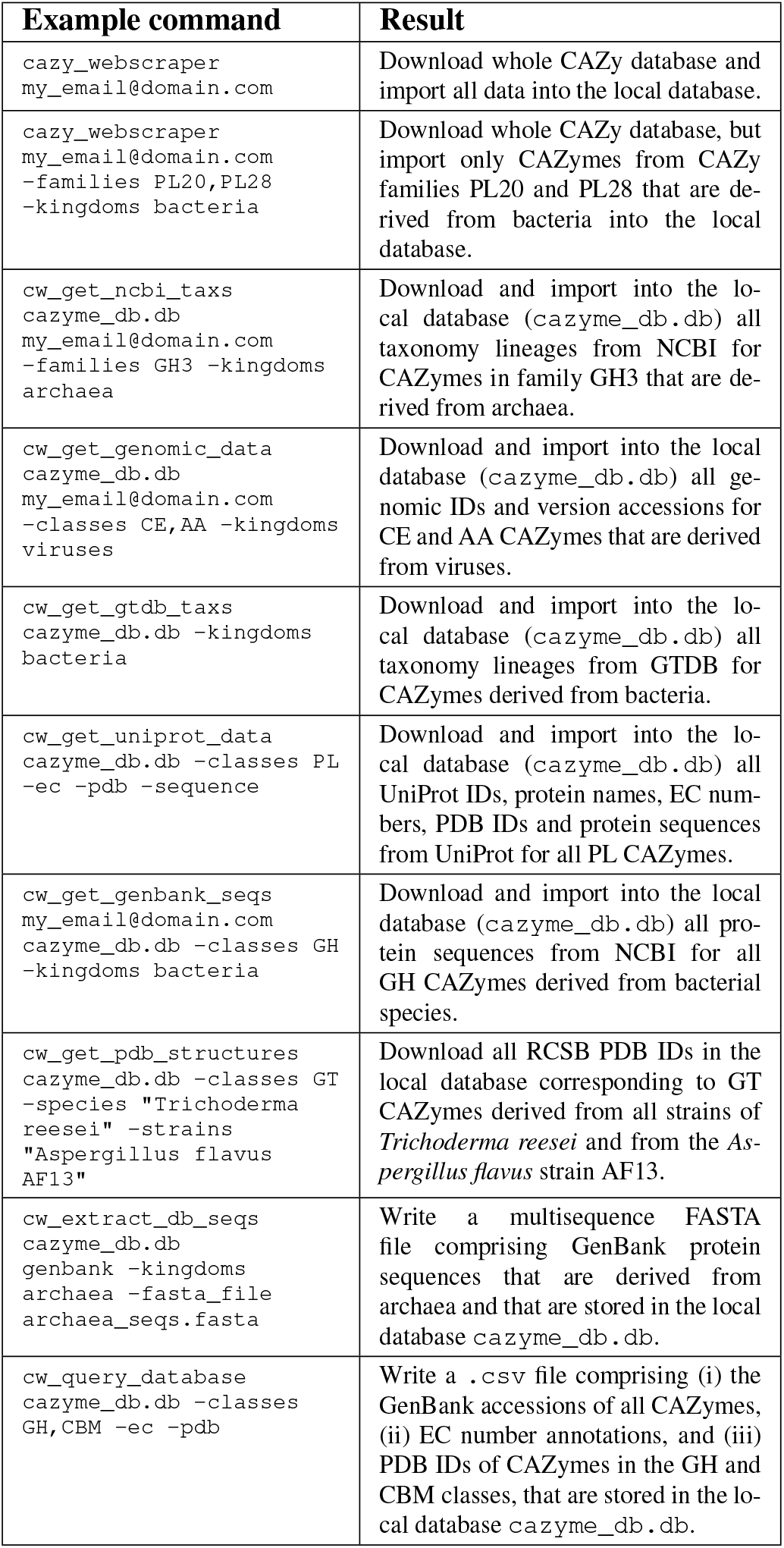
Example commands for importing data from the CAZy, UniProt, NCBI GenBank and RCSB PDB databases, and for data export.

### Automated retrieval of additional protein data, genomic data, sequences and structures

`cazy_webscraper` can extend the local CAZy database to incorporate protein sequences, functional annotations and structural data. `cazy_webscraper` uses NCBI accessions for each CAZyme to retrieve corresponding genome IDs, protein sequences, and other data using Biopython (15) and Entrez (16). UniProt data (UniProt ID, protein name, protein sequence, PDB ID, EC number, etc.) is retrieved using the BioServices package (17).

`cazy_webscraper` can update local CAZyme protein sequences if it detects a more recent version available from UniProt or NCBI. Protein sequences can be exported from the local database to FASTA format, or to a BLAST+ database (using the `--blastdb` flag).

PDB IDs obtained from UniProt can be used to obtain protein structures from RCSB PDB, using Biopython (18). PDB structure files are not stored directly in the database, but are saved to a user-specified location in the user’s preferred format (mmCIF, pdb, xml, mmtf or bundle). Database storage of structural data is planned in the `cazy_webscraper` development roadmap.

### Automated retrieval of NCBI and GTDB taxonomic classifications

`cazy_webscraper` uses the NCBI accession for each CAZyme record to retrieve and import the current taxonomic assignment directly from the NCBI Taxonomy (19) and/or GTDB databases (12). To link successfully with GTDB, a suitable NCBI genome assembly must be associated with the corresponding CAZyme accession.

### Querying the local CAZyme database

`cazy_webscraper` provides a command-line interface for common queries, returning output in CSV and/or JSON format. By default, the program returns only the NCBI accession for each CAZyme matching the provided criteria. The `--include` option allows reporting of additional fields, including CAZy class, CAZy family, CAZy subfamily, taxonomic kingdom, genus, host organism, GenBank protein sequence, UniProt accession, protein name, EC number, PDB accessions, and UniProt protein sequence.

The database generated by `cazy_webscraper` can also be queried directly by advanced users, using the SQLite console.

### Reproducible and shareable datasets and documentation

`cazy_webscraper` logs all data retrievals in the local database. The single compact database generated is time-stamped and shareable, facilitating reproduction of the downstream analyses that use it.

Data retrieval can be configured using a YAML file for precise reproduction. Reproducible local reconstruction of the database is assisted by caching unprocessed data from each data source (CAZy, NCBI, etc.). Each release of `cazy_webscraper` is associated with a unique digital object identifier (DOI), assigned by Zenodo.

### Installation

`cazy_webscraper` can be installed using Bioconda (20) or PyPI (21) package managers, or from source code (https://github.com/HobnobMancer/cazy_webscraper). `cazy_webscraper` is released under the MIT open licence.

## Results

### Performance

The complete CAZy database was downloaded, parsed, the data compiled into a local SQLite3 database, and taxonomic lineages recovered from the NCBI taxonomy database using `cazy_webscraper` in 14 min 53 s (±21 s). Excluding duplicate records, 2,232,090 unique CAZyme records were recovered. The resulting database was 270 MB in size and comprised proteins representing 199,076 taxa from five kingdom-level groups (bacteria, eukaryota, viruses, archaea and unclassified), spanning 688 CAZy families. A total of 108 records in the original CAZy database were found to be annotated with multiple inconsistent taxonomic lineages.

### Taxonomic distribution of CAZymes in CAZy

The CAZy database server does not provide summary information describing the total number of unique CAZymes, or the total number of CAZymes, in each taxonomic group (e.g. domain or kingdom). We retrieved the total number of CAZymes per taxonomic kingdom (archaea, bacteria, eukaryotes, viruses and unclassified) for each CAZy class as described in Methods (figure 2, time taken: 2 min 1 s, ±3 s).

**Fig. 2.**
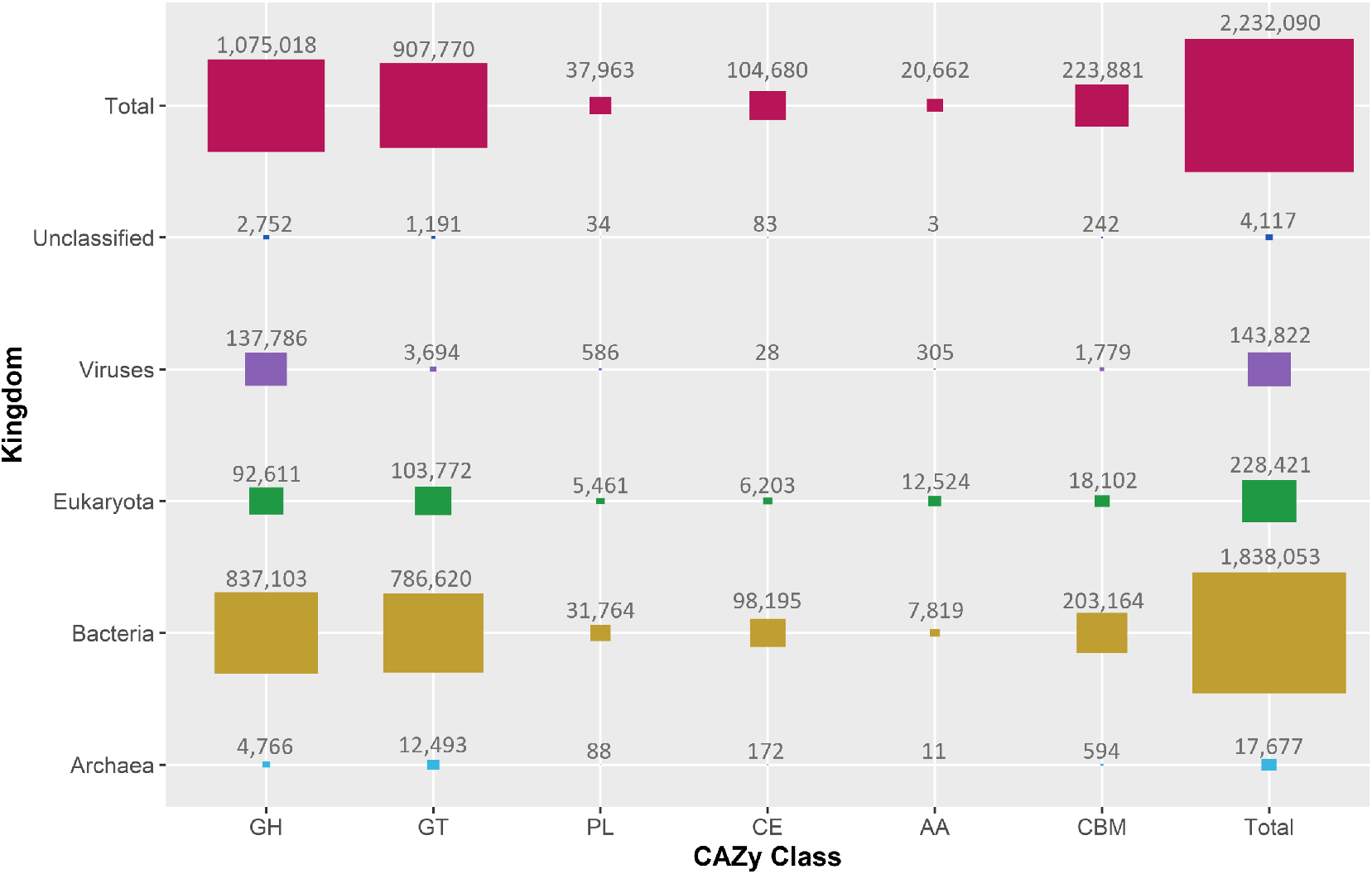
Proportional areas plot of the count of CAZymes in each class (in total, and by taxonomic kingdom), as recorded in CAZy (January 2022). Numbers show the count of unique sequence IDs associated with each combination of kingdom and CAZy class. Note that a single CAZyme may be assigned to several classes.

### CAZyme distribution by kingdom differs between CAZy and NCBI

As all CAZy records are drawn from NCBI, we expected that NCBI’s underlying sampling bias by taxonomic Kingdom, such as an overrepresentation of bacteria, might also be evident in CAZy. However, we found a statistically significant difference between the distributions of sequences across each kingdom in CAZy and in NCBI (*χ*^2^ test, p- value < 2.2E-16). We conclude that kingdom-level bias in CAZy does not simply reflect an underlying sampling bias in NCBI, but may represent a kingdom-level difference in relative abundance of CAZymes. We find specifically that “unclassified”, eukaryotic and archaeal sequences are relatively under-represented in CAZy, compared to what would be expected from the corresponding protein abundance in NCBI, but bacteria and viruses are over-represented (`additional file 1`).

### Relative and absolute representation of CAZymes varies by kingdom in CAZy

We wished to investigate how the relative abundance of CAZymes by species varies for each kingdom in CAZy. We expect this to represent both the underlying sampling bias across kingdoms and the differing prevalences of CAZymes in each kingdom. Table 2 presents kingdom-wise counts of: all species-level taxa in NCBI; the number of those taxa with at least one annotated CAZyme in CAZy; and the number of those taxa with at least 50 annotated CAZymes in CAZy. We find that less than 1.5% (29,283) of all species in the NCBI Taxonomy database have any CAZymes represented in CAZy, and less than 0.5% (6,693) of all species have more than 50 representative CAZymes in CAZy. Eukaryotic species are especially under-represented in the CAZy database.

**Table 2.**
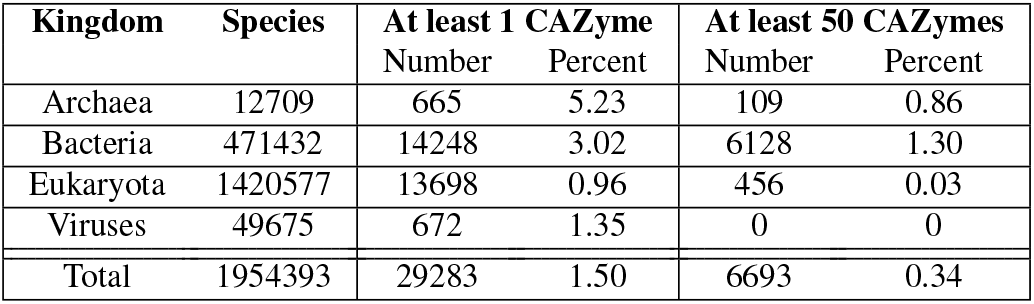
Counts of species-level taxa in NCBI with at least one, and with at least 50, CAZymes represented in CAZy. The percentage of all species in the corresponding kingdom that this represents is also shown.

| Kingdom | Species | At least 1 CAZyme |  | At least 50 CAZymes |  |
| --- | --- | --- | --- | --- | --- |
|  |  | Number | Percent | Number | Percent |
| Archaea | 12709 | 665 | 5.23 | 109 | 0.86 |
| Bacteria | 471432 | 14248 | 3.02 | 6128 | 1.30 |
| Eukaryota | 1420577 | 13698 | 0.96 | 456 | 0.03 |
| Viruses | 49675 | 672 | 1.35 | 0 | 0 |
| Total | 1954393 | 29283 | 1.50 | 6693 | 0.34 |

We find that the abundance of CAZy records within a species also varies by kingdom. Forty percent of bacterial species that have at least one CAZy record have more than 50 CAZy records, whereas this proportion falls to 16% for archaea, and 3% for eukaryotes. No virus species is associated with more than 50 CAZy records, likely because of their restricted genome size. We interpret these data potentially to be indicative of the differential expansion of CAZyme families within kingdoms.

### Auxiliary Activity (AA) families are expanded in eukaryotes and dominated by eukaryotic sequences

Most records in CAZy (82.35%) are bacterial CAZymes (figure 2). If CAZy classes were distributed evenly across all kingdoms, we would expect bacterial proteins to dominate in all cases. However, Auxiliary Activity (AA) CAZymes, including ligninolytic and lytic polysaccharide mono-oxygenases (6), are predominantly observed in eukaryotes. Seventeen of the 20 CE families, and 41 of the 43 PL families, are dominated by bacterial CAZymes. By contrast, 17 of the 18 AA families are dominated by eukaryotic enzymes, with several families only being represented in eukaryotes. We interpret this to suggest eukaryote-specific expansion of AA families, perhaps uniquely among CAZy classes. Kingdom-level distributions of all AA, CE and PL families are plotted in figure 3, and kingdom-level distributions for the complete set of CAZy families are plotted in `additional file 2`.

**Fig. 3.**
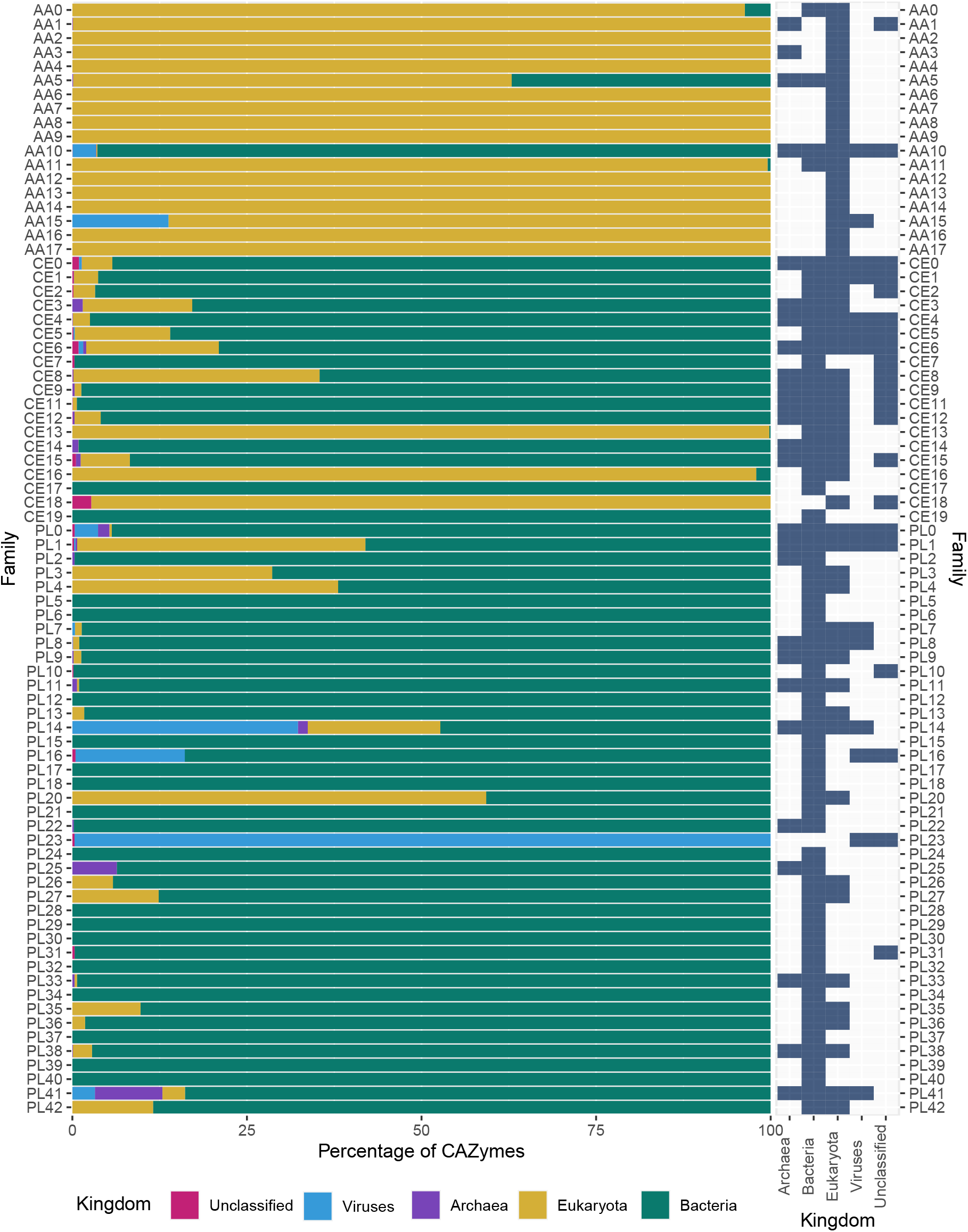
The proportion of CAZymes in each taxonomic kingdom for AA, CE and PL families (left), and absolute presence (blue)/absence (grey) in each kingdom (right). Families CE10 and PL19 are excluded as they have been withdrawn from the CAZy database.

Extending the comparison to all CAZy families (`additional file 2`), we find that 71.99% of all families (329) are dominated by bacteria, but only 22.98% (105) of families contain a majority of eukaryotic sequences. We note that, while CAZy families dominated by bacterial sequences tend to contain sequences from all other kingdoms (eukaryotes, archaea, viruses, unclassified), those families dominated by eukaryotic sequences have a different distribution (*χ*^2^ test, p-value = 4.517E-15) and a tendency to contain only eukaryotic sequences. We believe that this may be consistent with relatively recent expansion of CAZyme activities in eukaryotes, possibly in functions that are specifically beneficial for eukaryotic lifestyles, such as synthesis and degradation of chitin, lignin, and cellulose.

### Identifying under-represented archaeal groups in CAZy

CAZy catalogues taxonomic kingdom, genus, and species (and sometimes the strain) of each source organism, and further taxonomic detail is provided for each CAZy family by means of a Krona plot on the CAZy webserver. `cazy_webscraper` can retrieve the complete and most recent lineage from either or both of the NCBI Taxonomy and GTDB databases for all CAZymes in the local database, facilitating updated annotations, and analyses at all levels of phylogeny, across arbitrary sets of CAZymes for which a source genome can be identified.

To demonstrate this extended capability we summarise visually the complete taxonomic representation of all archaeal CAZymes in CAZy as an alluvial plot (figure 4). We exclude proteins belonging to organisms assigned *Candidatus* status, or that are catalogued in CAZy but no longer present in NCBI due to withdrawal or suppression of records. This and similar analyses to identify differential representation of arbitrary taxa in CAZy cannot currently be performed *via* the CAZy web service. A similar dataset for all archaeal CAZymes (including incomplete lineages and organisms assigned *Candidatus* status) reporting to genus level, is provided in `additional file 3`, again excluding proteins that are catalogued in CAZy, but no longer present in NCBI due to record withdrawal or suppression.

**Fig. 4.**
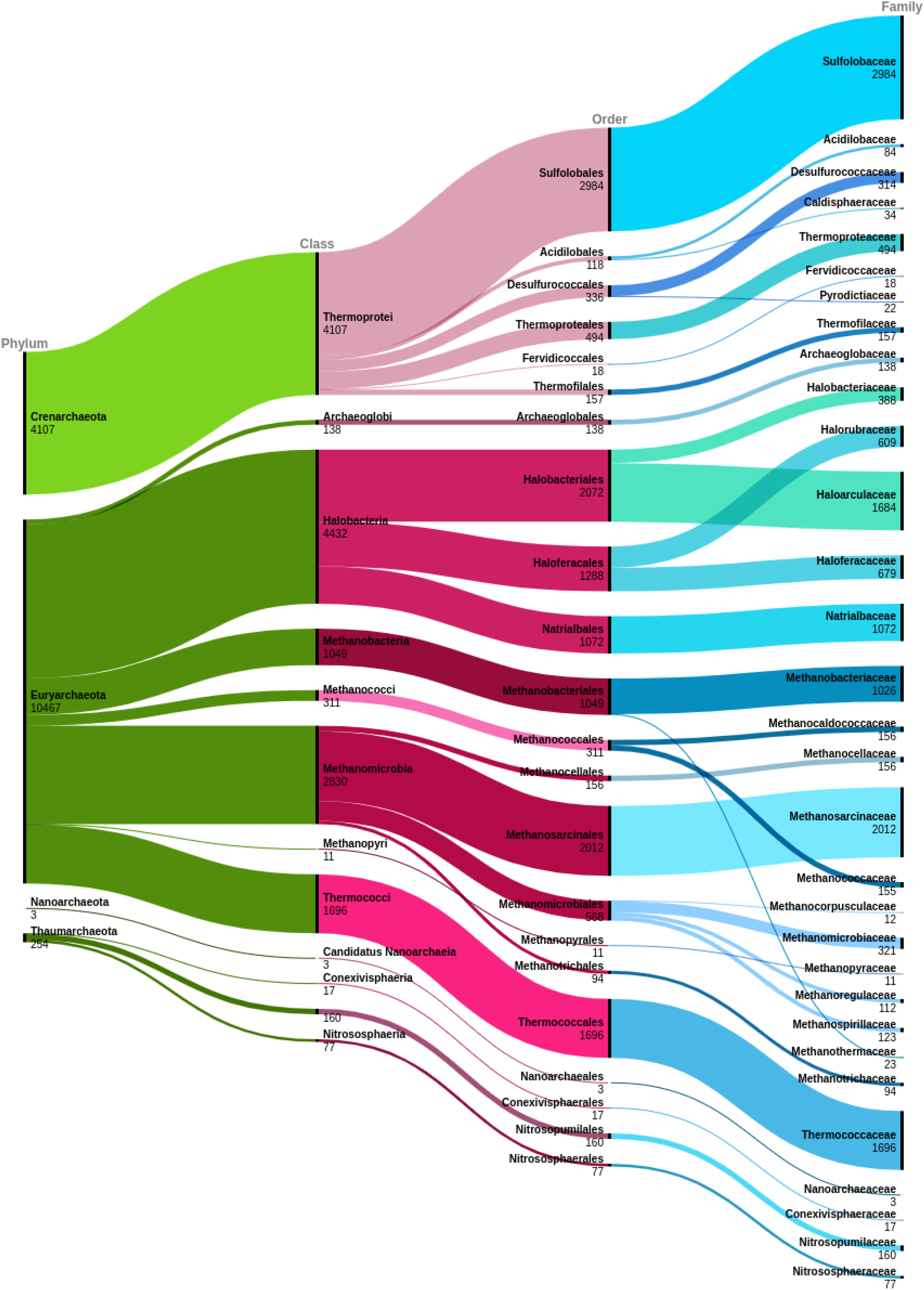
Alluvial diagram showing the count of CAZy records at each level of the NCBI archaeal taxonomy, from phylum to family. Incomplete (including “artificial sequences” and “environmental sample”) and *Candidatus* lineages are excluded.

By cross-referencing these data against the number of CAZymes listed per archaeal species name, we find that most archaeal CAZymes in CAZy derive from the phyla Euryachaeota and Crenarchaeota. Other archaeal phyla appear to be under-represented in CAZy, especially the Nanoarchaeota phylum which, despite over 250 sequenced genomes being available (May 2022), is represented by only 6 CAZymes in CAZy. Across the Archaea kingdom as a whole, at least one CAZyme was listed for every Archaeal family in the NCBI Taxonomy database, except for Conexivisphaeraceae and Nanoarchaeaceae. However, 35.26% of Archaeal genera in the NCBI taxonomy were not represented in CAZy.

We find that CAZyme coverage within archaeal lineages is also uneven. Within Crenarchaeota, the majority of CAZymes (approximately 75%) derive from the Sulfolobaceae family. The Thermoproteales, Thermofilales and Desulfurococcales orders are represented in CAZy ap-proximately proportionately to the number of corresponding genomes available at NCBI. However, assuming that 1-5% of coding sequences in a genome encode CAZymes (22), we find that the remaining orders Acidilobales (27 assemblies) and Fervidiocaccales (nine assemblies) are underrepresented, having only 81 and 18 CAZymes listed in CAZy, respectively. These orders may represent an opportunity for CAZyme mining.

### The PL20 CAZy family contains significant sequence diversity

Polysaccharide lyases (PLs) catalyse non-hydrolytic cleavage of glycosidic bonds and are exploited industrially for degrading biomass, for example in biofuel production (6, 23). CAZy family PL20 comprises 54 *β*-1,4-glucuronan lyases (EC 4.2.2.14), 22 from bacteria and 32 from eukaryotes, that cleave a *β*-1,4 linkage in the water-soluble homopolysaccharide polyglucuronate through *β*-elimination (24, 25). Only one member of the PL20 family, glucuronan lyase A (*gluc ly A*) from *Trichoderma reesei* NBRC 31329 (BAG80639.1), has been functionally and structurally characterised in the literature - to our knowledge - by January 2022. This single characterised representative is the main basis for functional annotation transfer on the grounds of common membership of PL20.

We explored sequence diversity within PL20 to investigate sequence similarity within the family, and to assess to what extent *gluc ly A* was likely to be representative of all PL20 members. Clustering of all PL20 protein sequences, as described in Methods, identified seven pairs of redundant sequences (figure 5), including GenBank record CEF86689.1 and its corresponding RefSeq sequence XP_391536.1. This indicates that the number of unique sequences in a CAZy family is not always the same as the number of sequence accessions.

**Fig. 5.**
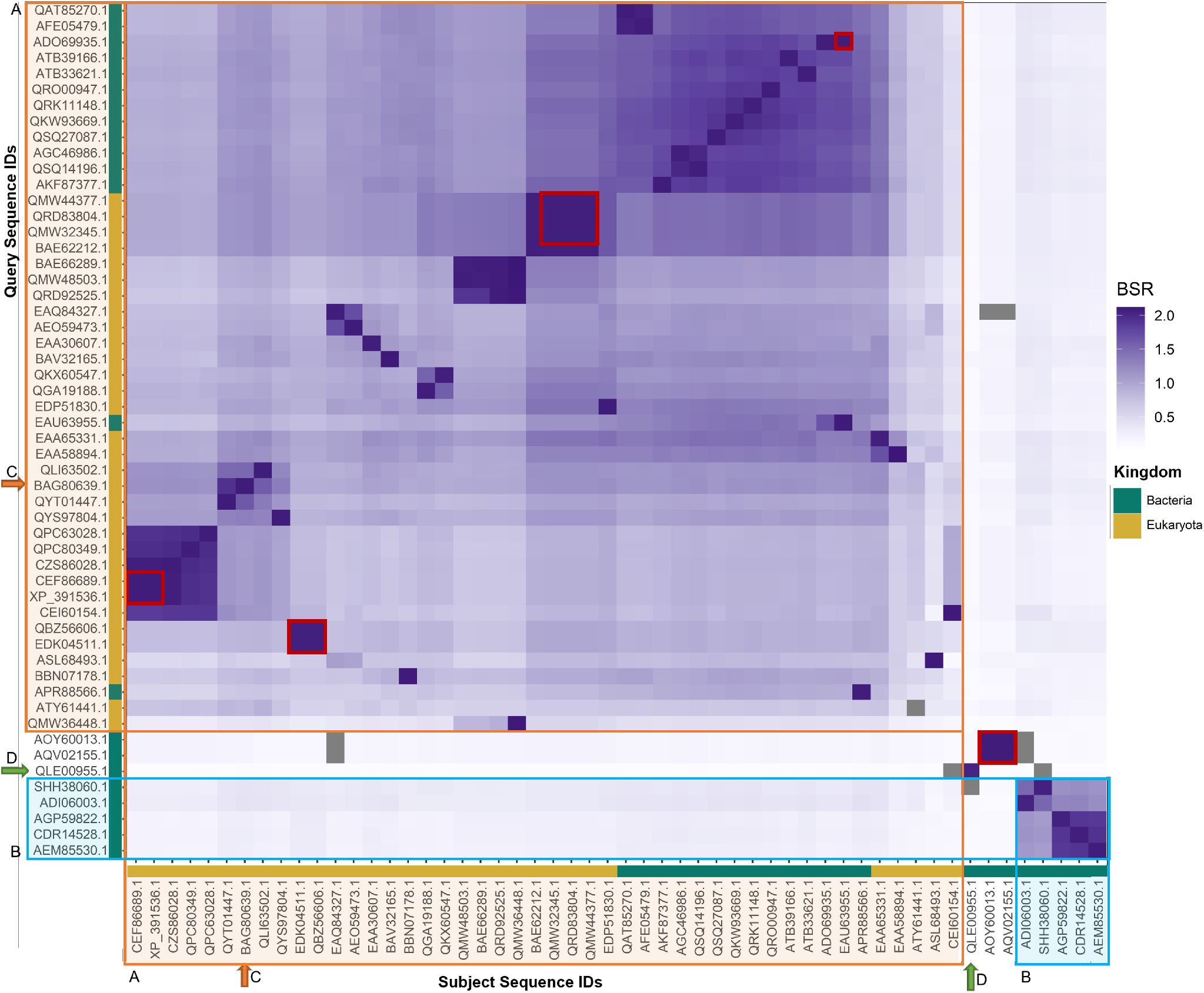
A heatmap representing BLASTP+ Score Ratios (BSRs) for CAZy family PL20 protein sequence pairwise alignments; colour intensity is proportional to BSR. The axis colour bars indicate bacterial (green) and eukaryote (gold) CAZymes. An apparently canonical glucuronan lyase group is indicated as group A (orange outline). Group B (blue outline) contains only PL20 sequences derived from Streptomycetes. The only protein to be functionally and structurally characterised in the literature is glucuronan lyase A from *Trichoderma reesei* (BAG80639.1), which is indicated with an orange arrow [C]. The glucuronan lyase A from *Galbibacter* sp. BG1 is indicated by the green arrow [D]. Redundant protein sequences are outlined in red.

Figure 5 shows a clustering of PL20 sequences on the basis of sequence similarity as measured by BLASTP+ Score Ratio (BSR). Four distinct clusters are observed, where the members of any one PL20 cluster share less than 40% pairwise protein sequence identity with members of any other cluster. This level of pairwise identity falls within or below the “Twilight Zone” of protein sequence identity that is a heuristic threshold for the ability to infer common structure or function on the basis of sequence identity (26). This raises the question of whether PL20 in fact comprises a functionally- or structurally-consistent family of proteins.

The largest, “canonical” group is a set of sequence-similar bacterial and eukaryotic CAZymes. This set includes *gluc ly A*, and shares an average of 55.4% pairwise amino acid identity across most of their length (±14.4% standard deviation, and mean 87.0% coverage ±23.0%) (group A, figure 5).

A second group comprises eight bacterial sequences (group B, figure 5) that share an average of only 28.7% pairwise identity (± 12.1% standard deviation, mean 79.0% ± 18.4% coverage) with the members of the “canonical” group. Group B contains all PL20 CAZy records deriving from *Streptomyces*. This group can be further subdivided into a pair of sequences from *Streptomyces sp. 3214*.*6* and *S. bingchenggensis BCW-1*, and a group of three sequences from *S. rapamycinicus NRRL 5491, S. iranensis* and *S. violaceusniger Tu 4113*.

The protein *HX109* (*HX109_05010*, GenBank QLE00955.1) from *Galbibacter* sp. BG1 (green arrow, figure 5) is relatively isolated within the heatmap, and shares low pairwise identity with all other PL20 sequences (BSR of less than 0.5, mean 23.8% ±7.77% identity and 34.0% ±20.6% coverage).

A redundant pair of sequences from *Desulfococcus multivorans* (sequence IDs AOY60013.1 and AQV02155.1) also have low pairwise identity with all other PL20 sequences (28.8% ±6.3% identity and 52.9% ±22.8% coverage). *gluc ly A* aligns only to the C-terminal region of AOY60013.1, suggesting the possible presence of an additional structural or functional domain comprising residues 1-175. However, dbCAN predicts no CAZyme domains in AOY60013.1 (`additional file 4`), and a BLASTP+ query against the NCBI nr database (`additional file 5`, accessed September 2022) matched this sequence to a conserved lignate_lyase2 superfamily domain (to which *β*-1,4-glucuronan lyases belong) across the full length of the AOY60013.1 sequence.

### A candidate membrane-associated PL20 CAZyme

To explore whether *HX109*, which also shares low pairwise sequence identity with all other PL20 family members, might share a similar fold with *gluc ly A* despite being below the usual “Twilight Zone” threshold for structural similarity (21.29% pairwise sequence identity), we superimposed the *gluc ly A* structure (PDB:2ZZJ) (25) onto the structure predicted by AlphaFold for *HX109*. The AlphaFold-predicted *HX109* structure possesses a fold similar to the canonical PL20 fold of *gluc ly A*: a *β*-jelly roll fold formed from two short *α*-helices and two antiparallel *β*-sheets (figure 6[A]). However, the calcium ion and citric acid binding residues in *gluc ly A* were not conserved in the predicted *HX109* structure (figures SI.1 and SI.2 in `additional file 6`). The optimal structural alignment obtained by superimposing the PDB:2ZZJ structure onto *HX109* PL20 domain residues 338-515 has a RMSD of 1.92 Å over 137 alpha carbons (C*α*).

**Fig. 6.**
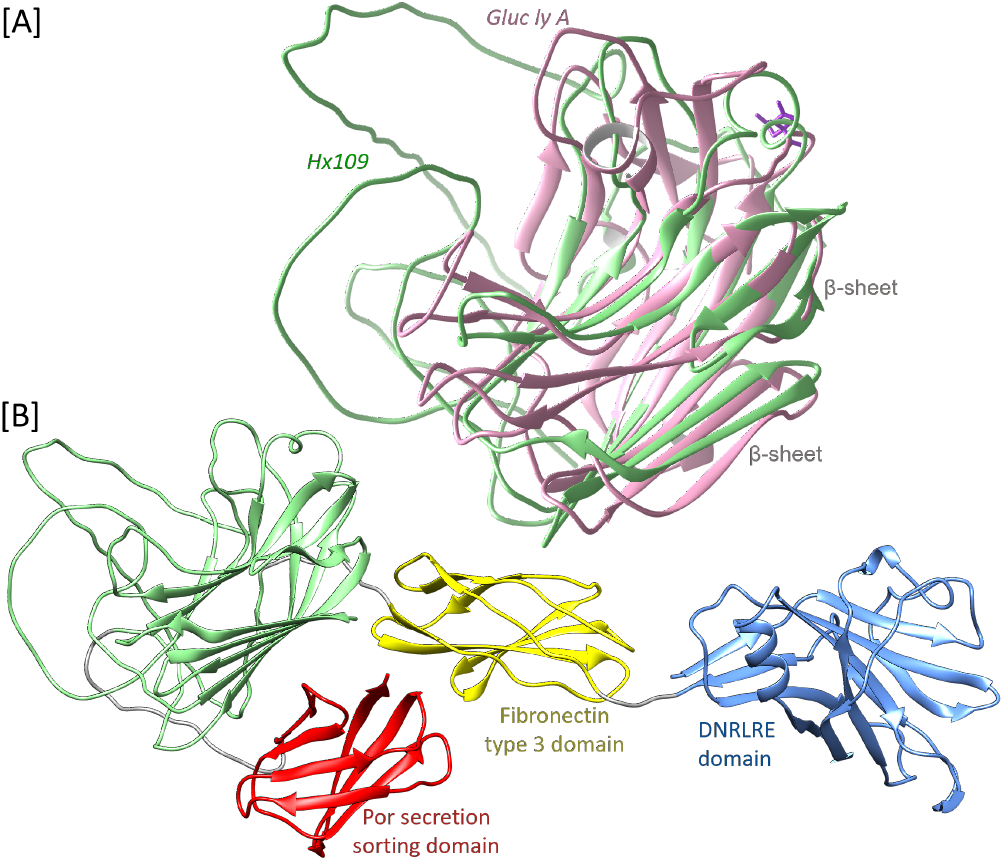
[A] The structure of *HX109* as predicted by AlphaFold (green), with the structure of *gluc ly A* (PDB:2ZZJ; pink) superimposed onto residues 338-515 (RMSD of 1.92 Å over 137 C*α*). [B] The complete predicted structure of *HX109*, which comprises from the Nto C-termini: DNRLRE domain (blue), fibronectin type 3 domain (yellow), PL20 domain (green), and the *por* secretion sorting domain (red). A citric acid molecule in PDB:2ZZJ is shown in purple.

The complete predicted structure of *HX109* is larger than a single PL20 domain. Instead, it appears to be composed of four domains, each with a pair of *β*-sheets. The global placement of each domain relative to the others is predicted with low confidence by AlphaFold (`additional file 7`) and so the complete conformation of the protein structure remains uncertain. However, a BLASTP+ query of the full length *HX109* protein sequence against the NCBI nr database (`additional file 8`) identified: (i) a conserved alginate lyase_2 superfamily (PFAM14099) domain that is also present in *gluc ly A*; (ii) a C-terminal *por* secretion sorting domain (Pfam:PF18962); and (iii) a fibronectin type 3 domain (Pfam:PF00041), which is found in many extracellular bacterial CAZymes (27, 28). A conserved N-terminal *DNRLRE* domain (NCBI Conserved Domains Database (CDD):NF033679), typically involved in cell wall function and structural organisation, was also identified (29). DeepTMHMM (30) did not predict any transmembrane domain in *HX109* (see Methods, `additional file 9`). SignalP predicted the presences of a signal sequence associated with a general secretion pathway. Taken together the signal peptide, along with the *por, fibronectin* and *DNRLRE* domains suggest that *HX109* may be secreted and perhaps a membrane-associated protein. The BLASTP+ query of *HX109* against all other CAZy PL20 sequences did not identify any of these indicators of potential membrane association. Therefore, *HX109* may be the only known candidate membrane-associated PL20 CAZyme.

### Identification of structurally-characterised carbohydrate esterases

Automated integration of sequence and structural data from databases such as UniProt and RCSB extends `cazy_webscraper`’s scope beyond simple replication of CAZy, and enables enhanced functional analysis of these proteins. To demonstrate this, we assessed the degree of structural characterisation of CE families by retrieving from UniProt all RCSB PDB IDs for the 104,680 CAZymes catalogued under the 20 CE families defined by CAZy. Obtaining this data took 3 mins 10 s (± 5.23 s, 3 replicates). The number of CAZymes in each CE family associated with at least one PDB ID was retrieved as described in the Methods (elapsed time: 15 mins 41 s, ±37 s, 3 replicates, table 3).

**Table 3.** The number of CAZymes in each CE family (January 2022), and the number of CAZymes in each family associated with at least one PDB ID in UniProt (April 2022).

| Family | Total CAZymes | CAZymes with PDB IDs in UniProt |
| --- | --- | --- |
| CE_0 | 2960 | 7 |
| CE_1 | 5026 | 6 |
| CE_2 | 709 | 4 |
| CE_3 | 600 | 3 |
| CE_4 | 34194 | 27 |
| CE_5 | 4195 | 10 |
| CE_6 | 453 | 1 |
| CE_7 | 2664 | 5 |
| CE_8 | 8927 | 7 |
| CE_9 | 19773 | 7 |
| CE_10 | 0 | 0 |
| CE_11 | 13842 | 4 |
| CE_12 | 3452 | 3 |
| CE_13 | 484 | 0 |
| CE_14 | 6487 | 6 |
| CE_15 | 583 | 8 |
| CE_16 | 189 | 0 |
| CE_17 | 18 | 1 |
| CE_18 | 37 | 1 |
| CE_19 | 251 | 1 |
| Total | 104844 | 101 |

Sixteen CE proteins that were not annotated as structurally characterised in CAZy were identified as having a solved structure by retrieving PDB IDs from their corresponding UniProt entry. However, thirty-one proteins annotated in CAZy as structurally characterised did not have an associated PDB ID recorded in their corresponding UniProt entry.

We find that CE families vary in the number of available representative structures. This information may guide strategic efforts to improve structural characterisation of families that are unrepresented or under-represented in the RSCB PDB. For example, no PDB IDs were retrieved for CE families 13 and 16, and only one CAZyme in each of the CE families 6, 17, 18 and 19 has been structurally characterised to date. These families may be worthwhile targets for structure determination.

### Identification of a possible novel CE domain variant

All members of a CAZy family are generally expected to share a common structural fold (31). CAZy families with no structural representatives may be the highest priority candidates for structural characterisation, but for protein sequences that share less than 40% identity it cannot be assumed that the backbone structure will be highly conserved (26). We have already established that there is significant sequence diversity in the PL20 CAZy family and, where a CAZy family can be subdivided into groups of proteins sharing more than 40% identity within, but less than 40% identity between, groups it may be that any such group having no experimentally-determined structural representative potentially possesses a structural fold that varies from the characteristic structure for the family as a whole. This represents a second, distinct group of strategic targets where structure determination may reveal novel folds.

To investigate potential structural variation in these low sequence identity groups, we used `cazy_webscraper` to retrieve all 251 protein sequences for CE19 family members from the NCBI Protein database. We clustered these sequences at a threshold of 40% identity and 80% coverage using `MMSeqs2`, and identified clusters containing no structurally characterised proteins (`additional file 10`). We noted that a predicted xylan esterase from *Luteitalea* sp. TBR-22 comprising 634 residues, *TBR22_41900* (*TBR22*, BCS34995.1), did not cluster with any other CE19 proteins by amino acid sequence. We used *TBR22* as a BLASTP+ query against the CE19 family, observing no match with amino acid identity above 32.17% (mean: 27.66%) (`additional file 11`). All such BLASTP+ alignments matched residues 1-380 in *TBR22*, implying the presence of a conserved N-terminal CE19 domain. However, few CE19 proteins matched against *TBR22* residues 380-634. This implied that *TBR22* contains a conserved CE19 domain (residues 1-380), and a second domain with unknown function (residues 380-634).

However, a single predicted acetylxylan esterase (*D6B99_08585*, AYD47656.1) was observed to align to *TBR22* residues 57-615, corresponding to residues 190-779 in *D6B99_08585*. dbCAN predicts that *D6B99_08585* residues 120-457 comprise a CE19 domain, but identifies no CAZy family domain for residues 458-779 (`additional file 12`).

dbCAN did not predict a CAZy family domain in *TBR22*. Querying the *TBR22* C-terminal domain (residues 380-634) against the NCBI nr database using BLASTP+ (see Methods) identified no putative conserved domains, but primarily returned hits against acetylxylan esterases from Acidobacteria (`additional file 13`). Using the sequence of *TBR22* residues 380-634 as a query against the PDB database also returned no matches (June 2022).

We predicted a candidate protein structure for *TBR22* using AlphaFold (`additional file 14`), and structurally superimposed the structure of PDB:6GOC (ALJ42174.1), the only CE19 structure listed in CAZy and UniProt, onto that prediction (figure 7[A]). 6GOC aligned well to the predicted CE19 domain (RMSD 2.05 Å across 311 C*α*), indicating that 6GOC and *TBR22* share a common *α*/*β* hydrolase fold, consisting of a three layer *α*/*β*/*α* sandwich containing a nine-stranded *β*-sheet. The two histidines in the 6GOC zinc ion binding site were conserved in the *TBR22* CE19 domain (figure SI.3 in `additional file 6`), suggesting that this function was retained. 6GOC was aligned onto the second *TBR22* domain (RMSD 2.24 Å across 200 C*α*) (figure 7[B]). The predicted second *TBR22* domain displays a *α*-*β*-*α* sandwich fold that resembles the 6GOC *α*/*β* hydrolase fold, but contains a seven-stranded, rather than a nine-stranded, *β*- sheet, and has two fewer *α*-helices below and one less *α*-helix above the *β*-sheet sandwich. Additionally, the zinc ion binding site in 6GOC was not conserved in the second *TBR22* domain (figure SI.4 in `additional file 6`). We note that the 6GOC structure excludes 190 N-terminal residues present in the full length ALJ42174.1 protein sequence. We cannot therefore conclusively assert that the second domain observed in *TBR22* is absent in the full length ALJ42174.1 sequence.

**Fig. 7.**
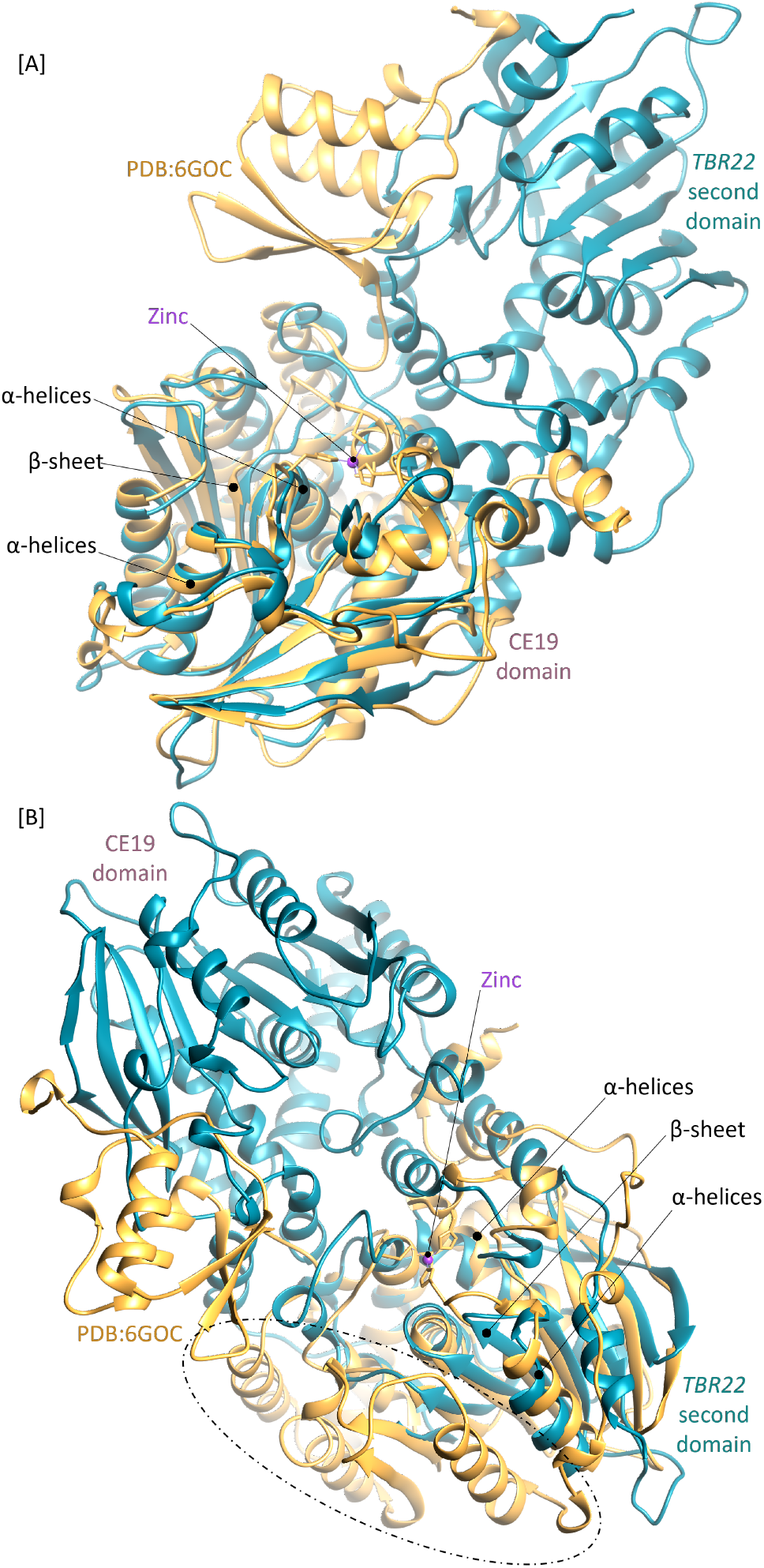
The structural fold of *TBR22_41900* (*TBR22*) as predicted by AlphaFold, shown by secondary structure in teal. Protein structure PDB:6GOC (shown in orange) superimposed onto [A] *TBR22* (RMSD 2.05 Å across 311 C*α*), and [B] onto the second *TBR22* domain (RMSD 2.24 Å across 200 C*α*) using the Chimera MatchMaker and Match-Align tools. The zinc ion in PDB:6GOC is shown in purple. Differences between 6GOC and the predicted *TBR22* fold are highlighted in the oval.

A BLASTP+ query of the *TBR22* C-terminal domain sequence against all CE protein sequences that have at least one associated PDB accession in UniProt (`additional file 15`) produced no alignment that matched more than 5% of the domain.

The full length *TBR22* sequence was queried iteratively against the RCSB PDB using HHPred to identify remote protein homologues not already catalogued in CAZy that may represent the full length and/or *TBR22* C-terminal domain structural fold in RCSB PDB (`additional file 16`). The three highest-scoring structures (probability score and E-value 99.5 to 99.7% and 4E-13 to 1.3E- 15 respectively): acetyl xylan esterase 3NUZ chain C (CAH07500.1); SusD/RagB-associated esterase-like protein 3G8Y (ABR41713.1); and RNA polymerase I subunit 6RUI chain A (AAA34992.1) were aligned against the full length *TBR22* predicted structure and against the *TBR22* second domain (see Methods) (`additional file 17`).

The structural alignments of 3NUZ and 3G8Y against full length *TBR22* (RMSD 1.85 Å across 275 C*α* and RMSD 1.95 Å across 278 C*α*, respectively) and the second *TBR22* domain (RMSD 1.89 Å across 209 C*α* and RMSD 1.97 Å across 215 C*α* respectively) are similar to those observed with 6GOC. 6RUI shows low structural similarity to full length *TBR22* (RMSD 2.23 Å across 231 C*α*) and the *TBR22* C-terminal domain (RMSD 2.35 Å across 174 C*α*) (`additional file 17`). We therefore propose that the second *TBR22* domain may potentially be a previously unobserved variant of the structural fold conserved in other CE19 enzymes.

### Prediction of a candidate novel CAZyme domain architecture - CE12:CBM35:PL11

We identified 174 clusters of CE12 protein sequences using `MMSeqs` (`additional file 18`). Only two of these clusters contained sequences with a structurally-characterised CE12 domain (We excluded ABN54336.1, which is associated with PDB ID 2W1W, as this structure represents only a CBM domain). We then attempted to determine if these two structurally characterised proteins (CAA61858.1, CAB15948.2) were likely to be representative for all CE12 family members. Eight of the 172 clusters having no characterised structure, comprising in total 50 sequences, contain proteins annotated with both CE12 and PL11_1 subfamily domains. Two enzyme in this group also include a CBM domain: QNU65479.1 contains a CBM35 domain and QZD56848.1 a CBM2 domain. To our knowledge none of the enzymes containing both CE12 and PL11_1 domains have yet been functionally and/or structurally characterised, so it was unclear to what degree the typical CE12 and PL11 activities, catalytic mechanisms and structures were conserved in these enzymes.

We arbitrarily chose the *Metabacillus* sp. KUDC1714 CAZyme *HUW50_16055* (*HUW50*, QNF30995.1) as a basis to explore whether a fold with CE12 and PL11 domains may already be structurally represented in the RCSB PDB database. The domain ranges of the CE12 and PL11 domains were annotated as described in Methods, which resulted in the unexpected prediction of an additional CBM35 domain in this protein (though we note that CBM35 domains are observed in other CE12 proteins). The CE12 domain was annotated as residues 22 to 440, the CBM35 domain as residues 440 to 644, and the PL11 domain as residues 806 to 1423. The structural fold for each domain was predicted independently using AlphaFold (`additional file 19`, figure 8).

**Fig. 8.**
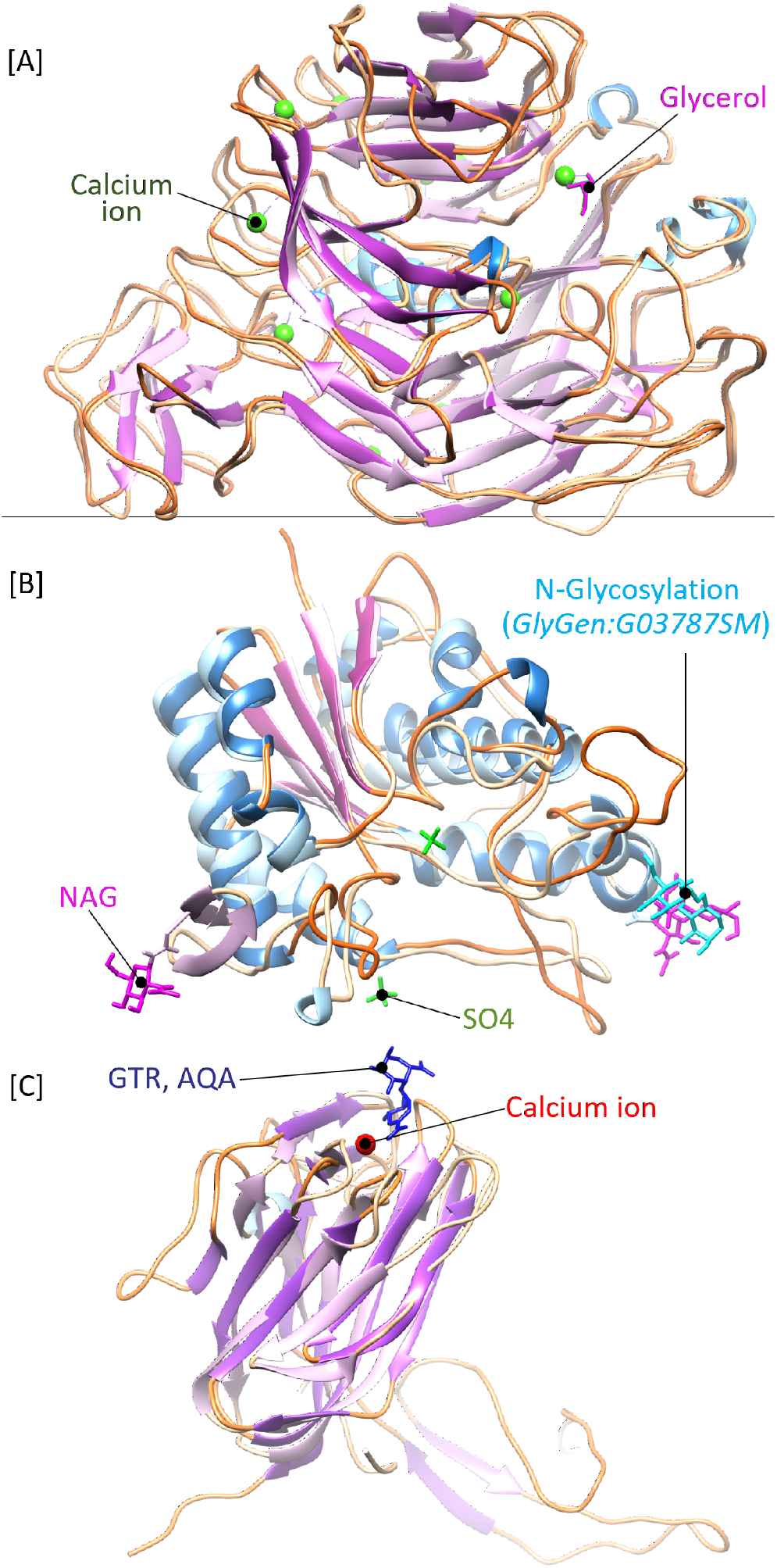
The AlphaFold-predicted structures for each of the three CAZyme domains of *HUW50_16055* (*HUW50*, NCBI QNF30995.1), showing the secondary structure. *α*-Helices are shown in blue, *β*-sheets in purple, the loop regions in orange, and calcium ions in dark green. The predicted structural folds are shown at greater opacity. [A] The predicted protein structure of *Huw50* PL11 domain superimposed onto PDB:4CAG with the 4CAG glycerol ligand shown in pink. [B] The predicted CE12 domain structure superimposed onto PDB:1DEO, with the 1DEO N-linked acetylglucosamine (NAG) shown in pink and the N-glycosylation (GlyGen:G03787SM) shown in light blue, and the two sulphate ions (SO4) shown in green. [C] The predicted CBM35 domain structure superimposed onto PDB:2W47 with the ligand 4-deoxy-*β*-L-threo-hex-4-enopyranuronic acid-(1,4)-*β*-D-galactopyranuronic acid (GTR, AQA) shown in dark blue.

The predicted PL11 domain structure in *HUW50_16055* forms a characteristic eight-bladed *β*-sheet propeller structure (figure 8[A]) (32–34), similar to the PL11 CAZyme RGL11 from *Bacillus licheniformis* (PDB: 4CAG) (RMSD 0.72 Å across 580 C*α*) (34). Additionally, the 10 calcium ion binding sites in PL11 CAZyme RGL11 are conserved in *HUW50* (figure SI.5 in `additional file 6`). The predicted fold of the *HUW50* CE12 domain forms the characteristic five-strand *β*-sheet sandwich between *α*-helices (35, 36) and aligns well to the CE12 enzyme RGAE from *Aspergillus oryzae* (PDB: 1DEO) (35) (RMSD 1.54 Å over 199 C*α*) (figure 8[B]). The RGAE catalytic triad, composed of a (nucleophilic) Ser, Asp and His, is conserved in the *HUW50* prediction (figure SI.6 in `additional file 6`), suggesting the enzyme may be catalytically active. However, the N-linked glycosylation site, comprising a short *α*-helix and small *β*-sheet in RGAE, was not predicted for *HUW50*. The CBM35 domain of *CTHE_3141* from *Acetivibrio thermocellus* (PDB: 2W47) aligns to the predicted structure of the *HUW50* CBM35 domain (RMSD 2.31 Å across 105 C*α*) (figure 8[C]), implying that *HUW50* does contain a CBM35 domain not yet annotated in its CAZy record. This highlights the difficulty of annotation for multi-domain CAZymes and, and that some classifications missed in the CAZy database may be identifiable if automated annotation were to be used.

## Discussion

CAZy is a valuable resource that provides a curated, sequence-based classification of carbohydrate processing enzymes. It enhances the community’s ability to develop predictors of enzyme function, catalytic mechanism, and structure, so is an ideal starting point for bioinformatic mining and cataloguing of the many carbohydrate-processing enzymes revealed by the rapidly-increasing corpus of publicly-available sequence data. Using CAZy for predictive tasks generally requires automated data processing, but the community’s ability to systematically use, or analyse, the data in CAZy is limited. This is in part due to the lack of an API for complex queries in the CAZy database, which is understandable given funding and resource constraints (6). Even with such an interface, users would need themselves to integrate CAZy data with sequence, structure, and other datasets, which presupposes some ability with data analysis and programming. `cazy_webscraper` provides a programmatic interface to the downloadable CAZy data, but also provides tools for analysing it that require no prior programming knowledge. This makes the dataset much more accessible and facilitates programming-free automated analysis for common tasks, including the use and extension of CAZy with data from public taxonomic, genomic and structure databases, with no additional burden on the CAZy resource beyond obtaining the publicly-available compressed database download. In this manuscript we demonstrate the use of `cazy_webscraper` to download the complete CAZy database, compile it into a local SQLite database, and extend it with additional information from public datasets, enabling reproducible interrogation of the CAZy database (table 1).

Download and analysis of the complete CAZy database with `cazy_webscraper` generates otherwise unavailable summary information about CAZy’s composition and comprehensiveness (figure 3) that are not necessarily evident when using CAZy’s web interface alone (figure 3). We find from our analysis that CAZy’s composition is influenced by the composition of its data sources. At a high level, NCBI contains many more bacterial sequences than eukaryotic sequences, and this is reflected in a similar distribution of CAZy records by kingdom. We also note that CAZy is simultaneously constrained by its data inclusion policies. For example, a maximum of 60 strains per species is considered for any CAZy family. This caps the sequence diversity that can be represented in the database (6), limiting the value of the dataset for within-species diversity analyses, and related pangenomic studies. By conducting large-scale predictions using tools such as dbCAN and incorporating the output into `cazy_webscraper`’s local database, these restricted datasets can be extended to facilitate such detailed studies.

Analysis of the complete CAZy dataset with `cazy_webscraper` enables data cleaning to refine the dataset for other purposes, such as generation of training sets for machine learning. For example, our survey of family PL20 demonstrates that CAZy contains redundant protein sequences, which might best be removed for some training sets. In particular, we identify cases where both GenBank and RefSeq accessions are catalogued as primary records for the same sequence, implying redundant representation of the same protein sequence by both accessions. Some RefSeq accessions included may represent Identical Protein Groups (IPGs), which have redundant amino acid sequences but may correspond to multiple proteins with distinct nucleotide coding sequences. Automated identification of such accessions is a benefit to phylogenetic reconstruction. We also show how CAZy summary data identifies opportunities for targeted functional and structural characterisation of under-studied groups of enzymes. This can assist more effective sampling, and expansion of our understanding of CAZyme diversity across taxa (figure 4).

CAZy does not store or directly provide sequence or structure data for its records. These data are instead linked from the CAZy record. This can place a non-trivial programming or network integration requirement on users of workflows employing CAZy in dynamic data integration with remote databases. `cazy_webscraper` provides commands for several actions simplifying this integration, including: (i) retrieving and locally storing sequence data for CAZy records; (ii) eliminating redundant records; (iii) allowing filtering and querying with user-defined criteria, to reduce the dataset to arbitrary groups of interest; and (iv) making filtered sequence sets available in standard formats for downstream alignment. `cazy_webscraper` facilitates integration of CAZy data with common bioinformatic analyses (clustering, phylogenetics, positive selection, CAZyme prediction, etc.).

We show that, by making sequence and structural data for CAZy records more conveniently available, `cazy_webscraper` helps identify sequences of strategic interest for further investigation. Such sequences may be potentially unrepresentative of their assigned CAZy classes or families, or otherwise of interest for functional or structural characterisation, potentially facilitating discovery of previously undisclosed CAZyme diversity (figure 5). In addition, this facilitates sequence-structure-function analyses such as mapping of conservation or correlated changes onto structure, potentially improving prediction of conserved sites and functional characterisation. In this manuscript we identify: (i) potentially the first example of a membrane-associated PL20 enzyme (from *Galbibacter*); (ii) a distinct cluster of *Streptomyces* PL20 sequences with minimal sequence similarity to other bacterial PL20s; (iii) a *Desulfococcus multivorans* protein with minimal similarity to any other PL20 record in CAZy; (iv) a potential new CE12 fold variant; and (v) a candidate novel CE12:CBM35:PL11 domain architecture. These preliminary analyses suggest that a wealth of novel CAZyme features may await discovery, with the ability to automatically interrogate the CAZy dataset.

### Extending CAZy’s taxonomic data

The CAZy database uses taxonomic assignments retrieved from NCBI. Unfortunately, taxonomic opinions are not always consistent between reference taxonomies, and may be revised over time within a single taxonomic resource (12). At NCBI taxonomic assignments are usually provided upon deposition by the sequence submitter but are not always updated in a timely manner to reflect revised understanding. This observation was a motivation for the widely-used Genome Taxonomy Database (GTDB) which aims to provide up-todate genome-based bacterial and archaeal taxonomies (12). `cazy_webscraper` can integrate assigned taxonomy from either or both of NCBI Taxonomy and GTDB, providing users with a choice and comparator of taxonomic references. For some downstream analyses, including building sequencebased CAZyme classifiers, it is desirable to account for bias in the input training sequence set, such as overrepresentation of a particular taxonomic group. We can identify likely influence of sampling bias in CAZy, but so as long as taxa are unevenly sampled in the underlying sequence databases from which CAZy is constructed it will remain difficult to rectify. Using `cazy_webscraper` we show that the overall distribution of CAZymes by kingdom reflects NCBI’s known sampling bias towards bacterial genomes (for example, the bacterium with the greatest number of RefSeq assemblies - 171,542 - is *Escherichia coli*, but the most-sequenced eukaryote, *Homo sapiens*, has only 1,268 (July 2022)). However, we find that CAZy class and family distribution at kingdom level is not only driven by NCBI database composition, but may also reflect biologically-significant causes. Specifically, we show that the AA class is dominated by eukaryotic sequences, consistent with the CAZy curators’ own observation that members of the AA class are strongly biased towards fungal enzymes (37) (figure 3), and also that many families of archaea are underrepresented in terms of their coverage in CAZy compared to NCBI (figure 4). `cazy_webscraper` provides a mechanism by which undersampled taxa may be identified, to help direct future strategic efforts in characterising CAZyme diversity. To facilitate this, an automated taxonomy survey of the CAZy dataset is in our development roadmap for `cazy_webscraper`.

### Extending structural information for CAZy records

We show how `cazy_webscraper` can extend the core CAZy dataset to include local annotation, sequence, taxonomic and structural data from NCBI, UniProt, GTDB and RCSB PDB databases, so it can be integrated more easily into extensive sequence, functional, structural and evolutionary studies. The CAZy website lists PDB IDs of CAZymes represented in the RCSB PDB database, but these must be identified and retrieved manually by the user. `cazy_webscraper` gathers PDB IDs from the corresponding UniProt record for each CAZyme. We find that identifying structural data *via* UniProt identifies additional PDB entries not currently listed in CAZy. For example, we find that nearly 20% of all CE class members associated with a specific PDB ID in UniProt are not annotated as structurally characterised in CAZy. We also find instances where CAZy lists PDB IDs for a record that are not annotated in the corresponding UniProt entry. We were thus able to use `cazy_webscraper` to consolidate and clean structural information provided in the CAZy dataset.

### Identifying potentially industrially-exploitable enzymes

CE enzymes remove methyl or acetyl groups that shield the polysaccharide backbone from degradation by GHs and PLs. Consequently, many CEs are industrially exploited to increase the surface area accessible to polysaccharide backbone-degrading enzymes, thus increasing the efficiency of polysaccharide degradation (38). A common substrate for enzymes for in several CE families is rhamnogalacturonan (RG), a structurally complex glycan that contains 13 unique monosaccharides and 21 distinct glycosidic linkages, nearly all of which require a bespoke enzyme for their cleavage (39). RG is highly abundant in plant cell walls and a rich source of monosaccharides for biofuel production (39, 40). It is possible that by mining CAZyme data it could be possible to extend the range of CE enzyme specificity available for industrial purposes, or to improve our understanding of sequence-structure-function relationships sufficiently to aid engineering of these enzymes.

Using `cazy_webscraper` we identified two CE enzymes having no representative structure in the PDB (*TBR22_41900* (*TBR22*) from *Luteitalea* sp. TBR-22, and a CE12-PL11 enzyme (with potentially a CBM35 domain) *HUW50_16055* (*HUW50*) from *Metabacillus* sp. KUDC1714). These enzymes shared relatively little (<40%) sequence identity with any other CE19 sequence in CAZy, and we suspected they might possess a novel or variant structural fold for the family. Their characterisation might, therefore, extend our coverage of the known CE19 protein structure space and aid elucidation of the enzyme family’s mechanism. Specifically, we identified a CE19 enzyme.

The CE19 family contains a single functionally characterised member, *BT1017* (AAO76124.1, (39)). *BT1017* is a pectin methylesterase that removes methyl esters from rhamnogalacturonan-II (RG-II) (39, 40). This activity facilitates complete degradation of the RG-II backbone by RGbackbone degrading enzymes (39). The structural fold shared by *BT1017* (PDB:6GOC) and the AlphaFold-predicted structure of the first domain of *TBR22* are similar, and it is possible that *TBR22* could possess the same RG-II-backbone degradation behaviour as *BT1017*. However, the predicted structural fold of the second *TBR22* domain did not show similarity with any structures in the RSCB PDB, which suggests it might play a different role or target an unknown substrate, perhaps working synergistically with the *TBR22* CE19 domain. This suggests that *TBR22* warrants further investigation and that CE19, and perhaps other CAZy families, harbour greater structural and functional diversity than is suspected.

Family CE12 comprises acetylesterases that target plant cell wall polysaccharides: pectin, RG and xylan (EC 3.1.1.-). Several characterised CE12 enzymes display synergistic activity resulting in an increased rate of polysaccharide degradation. For example, synergy between the RG-lyase *YesT* and a xylanase significantly increases the rate of degradation of acetylated xylan (41). CAZy family PL11 represents rhamnogalacturonan lyases (EC 4.2.2.23 and EC 4.2.2.24) that degrade the RG backbone via *β*-elimination (42). The synergistic activity between a RG-acetylesterase and the RG backbone degrading enzymes RGase A and B is known to significantly increase the rate of RG degradation (43). We also predict that *HUW50* contains a CBM35 domain, examples of which are known to target RG (44). We therefore speculate that the *HUW50* CE12-PL11 protein contains a PL11 domain that acts synergistically with CE12 in a novel, synergistically-linked domain composition to more efficiently degrade RG.

*HUW50* is only one example of 50 candidate multidomain CE12:PL11 proteins we identify for the first time in our study. No protein structures in the RCSB PDB represent the global structure of these enzymes. The synergistic activity we speculatively propose has not been directly experimentally determined, nor have their substrate(s) been fully characterised. However, the family activities listed in CAZy imply that these enzymes may degrade plant cell-wall polysaccharides, and that investigation of their properties could be of potential biotechnological interest.

## Conclusion

`cazy_webscraper` automates retrieval and integration of user-specified datasets from CAZy, NCBI, GTDB, UniProt, and RCSB PDB databases to create a local CAZyme database of proteomic, taxonomic, and structural data. The commands used to create the dataset can be provided in a YAML configuration file and shared or re-run for reproducible creation and update of datasets. The compilation of CAZy records into a comprehensively-logged SQLite3 database enhances transparency, reproducibility, shareability, and replicability of analyses. Thus, `cazy_webscraper` facilitates mining an extended CAZy dataset *via* bioinformatic discovery workflows, enabling identification of enzyme candidates of interest for novel functional and/or structural characterisation. We expect this to be of particular use for industrial biotechnology applications, and in particular biofuel production.

## Methods

`cazy_webscraper` version 2.0.13 (DOI:10.5281/zenodo.6343936) was used for all examples in this manuscript. All `bash` commands used to augment, query and extract data from the local CAZyme database are provided as executable bash scripts in `additional file 20`, alongside a README file walk-through.

Manuscript figures were generated using an RMarkdown notebook (`additional file 1` is an archive including this notebook and all data files required to run the analyses). All analyses were performed on a consumer-grade laptop with AMD FX-6300 (3.5 GHz) processor, 16 GB RAM, and an onboard SSD drive and theoretical network speed of 107 Mbps, running Ubuntu 22.04.2 LTS. All timings were measured in triplicate, and variation reported as standard deviation. For all SQL commands executed in the SQLite3 console the number of unique protein sequence accessions returned by the query was interpreted as the count of CAZymes matching the query. Default parameters were used for all software, unless otherwise stated.

### Building the local reference CAZyme database

The CAZy database was downloaded from http://www.cazy.org on 13th January 2022 and used to create a local database with `cazy_webscraper`, configured using the `build_database.sh` script (`additional file 20`).

### Taxonomic assignments in the local CAZyme database

During testing, we found that some CAZy database records were assigned to taxa with no corresponding entry in the NCBI Taxonomy database. `additional file 21` lists 108 proteins assigned to multiple source organisms in the July 2022 CAZy release).

The number of CAZymes per taxonomic kingdom as recorded in CAZy, and the count of CAZymes per taxonomic kingdom for each CAZy class and each CAZy family, were obtained with the script `cazymes_per_cazy_kingdom.sh` (`additional file 20`). Additional NCBI taxonomic assignments were imported using the script `get_ncbi_taxs.sh` (`additional file 20`).

The count of proteins in the NCBI GenBank database per kingdom was retrieved from the NCBI Taxonomy database (https://www.ncbi.nlm.nih.gov/taxonomy accessed May 2022). A *χ*^2^ test (`additional file 1`) was used to test whether the distribution of CAZymes per taxonomic kingdom recorded in CAZy differed from the distribution of all proteins in the NCBI protein database by kingdom. A matrix containing the explained variance per kingdom was calculated for this *χ*^2^ model (`additional file 1`):

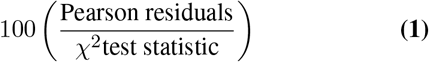

Pearson residuals are defined, where *O* represents observed and *E* represents expected values:

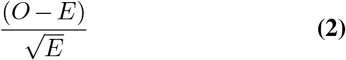

A second *χ*^2^ test was used to test whether the presence of CAZymes per taxonomic kingdom in CAZy families dominated by eukaryotes differed from CAZy families dominated by bacteria; a matrix containing the explained variance per kingdom was calculated for this *χ*^2^ model (`additional file 1`).

Lineage information and the count of CAZymes in each NCBI taxon were obtained for all archaeal CAZymes in the local database using the script `get_archaeal_cazymes.sh` (`additional file 20`). Incomplete lineages were manually removed from the data, as were all lineages assigned as *Candidatus*. `additional file 22` lists all archaeal NCBI lineages in the resulting local database. These remaining assignments were used to construct alluvial diagrams of the number of CAZymes at each node of the archaeon lineage (RAWGraphs version 2.0, (45)).

### Survey of sequence diversity in the PL20 family

Protein sequences of CAZymes in family PL20 were downloaded from NCBI and added to the local CAZyme database using `cazy_webscraper` (`get_pl20_seqs.sh`, `additional file 20`). The sequences were extracted to a multi-sequence FASTA file (`additional file 23`). All-vs-all pairwise sequence comparison was performed with these sequences using BLASTP+ v2.13.0 (46); configured using script `blastp_pl20.sh` in `additional file 20`. Corresponding taxon assignments were extracted from the local database (`get_pl20_taxons.sh`, `additional file 20`). Using `additional file 1`, pairwise sequence similarity across the family was calculated as the BLAST Score Ratio (BSR) (47) for each top-ranked pairwise alignment, and the R package `heatmaply` v1.3.0 (48) used to cluster the resulting matrix (`euclidean` distance, `hclust` function). Bar charts were produced plotting the taxonomic kingdom distribution across each axis of the heatmap using `ggplot2` v3.3.5 (49).

The *HX109_05010* (*HX109*, GenBank accession QLE00955.1) protein sequence was queried using BLASTP+ v2.13.0, with the BLOSUM45 matrix against the NCBI non-redundant (nr) protein database (accessed 5th May 2022), to identify candidate conserved functional domains (`additional file 8`).

### Prediction of signal peptide and transmembrane domains

Signal peptides were predicted using `signalP` (version 6.0, (50)). Transmembrane domains were predicted using `DeepTMHMM`, via the DeepTMHMM server (https://dtu.biolib.com/DeepTMHMM) (30)).

### Structural fold prediction and structure superimposition

Structural folds were predicted for individual domains of proteins (excluding signal peptides) using the official AlphaFold Colab notebook (version 2.1.0, (51)). Structures obtained from RCSB-PDB were superimposed onto the predicted structures and Root Mean Square Distance (RMSD) calculated using the `MatchMaker` tool, followed by the `Match-Align` tool, in UCSF Chimera (version 1.16, (52, 53)).

### Structural survey of CE families

RSCB-PDB accessions for all CE class CAZymes in the local database were retrieved from UniProt (accessed April 2022). The number of CAZymes in each CE family with at least one PDB ID was obtained using the script `get_ce_pdbs.sh` (`additional file 20`).

Protein sequences for all CE family members were downloaded from NCBI, imported into the local database, and written to a multi-sequence FASTA file (`additional file 24` (CE19); `additional file 25` (CE12)) using `cazy_webscraper`, configured using using script `get_ce_seqs.sh` (`additional file 20`. These sequences were clustered with `MMseqs2` (v13.45111 (54)) with thresholds of 40% identity and 80% coverage (using `cluster_ce12_ce19.sh` in `additional file 20`). The resulting clusters containing proteins with corresponding structures deposited in RCSB-PDB were identified using the Python script `gather_clusters.py` in `additional file 20`.

### Functional and structural prediction for CE19 enzyme *TBR22_41900* (*TBR22*)

The full-length protein sequence of the CE19 CAZyme *TBR22_41900* (*TBR22*, GenBank ID BCS34995.1) was chosen to represent a cluster of CE19 proteins identified by `MMSeqs2` but having no structural representative in the RCSB-PDB. The protein sequence of *TBR22* was queried against all other CE19 family members in the local database using BLASTP+ (`blastp_ce19.sh`, `additional file 20`; results in `additional file 11`). CAZy family domains were predicted for this sequence and for *D6B99_08585* (AYD47656.1, `additional file 12`) using dbCAN v3.0.2 (55).

The C-terminal domain of *TBR22* (residues 380-634) was queried against the NCBI nr protein database using the NCBI BLASTP+ server (results in `additional file 13`), and against the RSCB-PDB database, using the PDB server (both accessed June 2022).

An iterative sequence-based search for structural homologues was made using the `HHpred` server (v2.1, https://toolkit.tuebingen.mpg.de/tools/hhpred, (56)) with the full-length *TBR22* protein as query to the RCSB-PDB database (`additional file 16`). The structural fold of *TBR22* was predicted using the AlphaFold Colab service (`additional file 14`). Structures obtained from the `HHpred` search were superimposed onto the predicted structure as described above (`additional file 17`).

Protein sequences for all CE enzymes with at least one representative PDB ID in the local CAZyme database were written to a multi-sequence FASTA file (`get_ce19_pdb_prots.py`, `additional file 20`). The C-terminal domain of *TBR22* was queried against these sequences using BLASTP+ to identify potential homologues (`blastp_cterm_ce19.sh`, `additional file 20`; results in `additional file 15`).

### Annotation and structural fold prediction of CAZyme domains in *HUW50_16055* (*HUW50*)

CAZyme domains were predicted for the protein *HUW50_16055* (*HUW50*, QNF30995.1, `additional file 26`) using dbCAN (version 3.0.4). The full-length sequence of *HUW50* was queried against the protein sequences of two structurally characterised CE12 CAZymes, three structurally characterised PL11 CAZymes, and 16 structurally characterised CBM35 CAZymes using BLASTP+ (`additional files 27-29`; queries against the CBM35 sequences used the BLOSUM45 matrix). *HUW50* CAZyme domains were annotated using dbCAN (`HMMer` results) and the BLASTP+ sequence alignments.

Structural folds were predicted separately for each predicted CAZyme domain in *HUW50* using AlphaFold Colab (`additional file 19`), and experimentally-derived structures superimposed onto the corresponding *HUW50* domain using UCSF Chimera as described above. Metal ion and substrate binding site annotations were manually retrieved from UniProt.

## Funding

E.E.M.H. is funded by a BBSRC EASTBIO Doctoral Training Partnership award.

## Abbreviations

AA: Auxiliary activity
BSR: BLAST Score Ratio
CBM: Carbohydrate binding module
CE: Carbohydrate esterase
EC: Enzyme Commission
GH: Glycoside hydrolase
GT: Glycosyl transferase
LPMO: Lytic polysaccharide monooxygenase
NCBI: National Centre for Biotechnology Information
nr: NCBI non-redundant database
ORM: Object relationship model
PDB: Protein Data Bank
PL: Polysaccharide lyase
RCSB: Research Collaboratory for Structural Bioinformatics
SQL: Structured Query Language.

## Availability of data and materials

Project name: `cazy_webscraper` Project home page: https://hobnobmancer.github.io/cazy_webscraper/ GitHub Repository: https://github.com/HobnobMancer/cazy_webscraper. Documentation: https://cazy-webscraper.readthedocs.io/. Operating systems: Linux, MacOS and Windows 7 or higher. Programming language: Python 3.9. License: MIT License. Any restrictions to use by non-academics: Commercial rights reserved.

## Competing interests

The authors declare that they have no competing interests.

## Authors’ contributions

EEMH wrote the software; EEMH, TMG, LP designed the software; EEMH, LP wrote documentation; EEMH, TMG, LP conceived the analyses; EEMH performed the analyses; EEMH, TMG, LP wrote, read and approved the final manuscript.

## Consent for publication

Not applicable.

## Ethics approval and consent to participate

Not applicable.

## Additional files

**Additional file 1 — RMarkdown notebook**. Zip file containing a RMarkdown notebook and all required data files to perform statistical tests and generate figures. (ZIP 58KB)

**Additional file 2 — Kingdom distribution per CAZy family**. The proportions and presence/absence plots of CAZymes per taxonomic kingdom in each CAZy family. (PDF 46KB)

**Additional file 3 — NCBI archaeal lineages**. Alluvial plot for all lineages retrieved from NCBI for all archaeal CAZymes in CAZy, from kingdom to genus. (SVG 175KB)

**Additional file 4 — AOY60013.1 dbCAN output**. dbCAN prediction of CAZyme domains in NCBI protein AOY60013.1. (ZIP 3KB)

**Additional file 5 — BLASTP AOY60013.1 vs NCBI nr**. BLASTP protein NCBI AOY60013.1 against the NCBI nr database (September 2022). (ZIP 157KB)

**Additional file 6 — Binding site structure comparison**. Additional annotations of conserved active site and binding site residues in *HX109_05010, TBR22_41900, HUW50_16055* and PDB 6GOC, 4CAG, 1DEO and 2W47. (PDF 1577KB)

**Additional file 7 — *HX109_05010* predicted structural fold**. All outputs from Alphafold (including MSA coverage and probability plots) for *HX109_05010*. (ZIP 383KB)

**Additional file 8 — BLASTP *HX109_05010* vs NCBI nr**. BLASTP protein *HX109_05010* against the NCBI nr database (May 2022). (CSV 13KB)

**Additional file 9 — DeepTMHMM *HX109_05010* output**. All outputs from DeepTMHMM for *HX109_05010*. (ZIP, 59KB).

**Additional file 10 — CE19 clusters**. CSV file summarising CE19 clustering, cluster members and proteins represented in RCSB PDB. (ZIP 3KB)

**Additional file 11 — BLASTP *TBR22_41900* vs CE19**. BLASTP *TBR22_41900* against CE19. Output from querying *TBR22_41900* against all CE19 members. (CSV 5KB)

**Additional file 12 — *D6B99_08585* dbCAN output**. Predicted CAZyme domain output from dbCAN for *D6B99_08585*, AYD47656.1. (ZIP 2KB)

**Additional file 13 — BLASTP *TBR22_41900* vs NCBI nr**. Output from BLASTP *TBR22_41900* against the NCBI nr database (June 2022). (CSV 25KB)

**Additional file 14 — *TBR22_41900* predicted structural fold**. All outputs from Alphafold (including MSA coverage and probability plots) for *TBR22_41900*. (ZIP 602KB)

**Additional file 15 — BLASTP *TBR22_41900* output**. BLASTP output of *TBR22_41900* queried against all characterised CE structures. (CSV 2KB)

**Additional file 16 — HHpred output**. Output from querying *TBR22_41900* against the PDB database using HHpred (June 2022). (ZIP 232KB)

**Additional file 17 — CE19 *TBR22_41900* structure comparison**. Result of superimposing PDB structures 6GOC, 3GY8, 3NUZ and 6RUI onto the predicted *TBR22_41900* structure. (DOCX 3758KB)

**Additional file 18 — CE12 clusters**. CSV files summarising CE12 clustering, cluster members and proteins represented in RCSB PDB. (CSV 44KB)

**Additional file 19 — Predicted *HUW50_16055* structural folds**. All outputs from Alphafold (including MSA coverage and probability scores) for *HUW50_16055* CAZyme domains. (ZIP 1181KB)

**Additional file 20 — Scripts to perform analyses**. ZIP file containing Bash and Python scripts used to perform the presented analyses, alongside a README file walk-through. (ZIP, 23KB).

**Additional file 21 — Proteins with multiple taxonomies**. Log file generated by `cazy_webscraper` listing all instances where multiple source organises were listed for the same protein. (TXT 15KB)

**Additional file 22 — Archaeal lineages**. Archaeal lineages. All lineages retrieved from NCBI Taxonomy for all archeal CAZymes in the local CAZyme database. (CSV 13KB)

**Additional file 23 — PL20 sequences**. FASTA file of protein sequences for all PL20 members. (FASTA 16KB)

**Additional file 24 — CE19 sequences**. FASTA file of protein sequences for all CE19 members. (FASTA 148KB)

**Additional file 25 — CE12 sequences**. FASTA file of protein sequences for all CE12 members. (FASTA 1493KB)

**Additional file 26 — dbCAN *HUW50_16055* output**. dbCAN CAZyme domain predictions for *HUW50_16055*. (ZIP 3KB)

**Additional file 27 — *HUW50_16055* CE12 alignment**. Alignment from querying *HUW50_16055* against structurally characterised CE12 proteins using BLASTP. (TXT 3KB)

**Additional file 28 — *HUW50_16055* PL11 alignment**. Alignment from querying *HUW50_16055* against structurally characterised PL11 proteins using BLASTP. (TXT 4KB)

**Additional file 29 — *HUW50_16055* CBM35 alignment**. Alignment from querying *HUW50_16055* against structurally characterised CBM35 proteins using BLASTP. (TXT 5KB)

## Notes

https://zenodo.org/record/7389830#.Y4nXypb7S70

### Competing Interest Statement

The authors have declared no competing interest.

## References

1. D. Chettri, A. K. Verma, and A. K Verma. Innovations in cazyme gene diversity and its modification for biorefinery applications. Biotechnology Reports, 28, 2020.

2. B. Zhang, Y. Gao, L. Zhang, and Y. Zhou. The plant cell wall: Biosynthesis, construction, and functions. Journal of Integrative Plant Biology, 63(1):251–272, 2016.

3. Y. Liu, R. Li, J. Wang, X. Zhang, R. Jia, Y. Gao, and H. Peng. Increased enzymatic hydrolysis of sugarcane bagasse by a novel glucose- and xylose-stimulated β-glucosidase from anoxybacillus flavithermus subsp. yunnanensis e13t. BMC Biochemistry, 18(1), 2017.

4. M.R. Kao, H.W. Kuo, C.C. Lee, K.Y. Huang, T.Y. Huang, C.W. Li, C.W. Chen, A.H.J. Wang, S.M. Yu, and T.H.D. Ho. Chaetomella raphigera β-glucosidase d2-bgl has intriguing structural features and a high substrate affinity that renders it an efficient cellulase supplement for lignocellulosic biomass hydrolysis. Biotechnology for Biofuels, 12(1):258, 2019.

5. M. Zheng, K. Zhang, J. Zhang, L. Zhu, G. Du, and R. Zheng. Cheap, high yield, and strong corn husk-based textile bio-fibers with low carbon footprint via green alkali retting-splicing-twisting strategy. Industrial Crops and Products, 188:115699, 2022.

6. E. Drula, M. Garron, S. Dogan, V. Lombard, B. Henrissat, and N. Terrapon. The carbohydrate-active enzyme database: functions and literature. Nucleic Acids Research, 50(D1):D571–D577, 2022.

7. R. V. Honorato. Cazy-parser a way to extract information from the carbohydrate-active enzymes database. The Journal of Open Source Software, 1(8), 2016.

8. Google. Bigquery, 2022.

9. E. W. Sayers, M. Cavanaugh, K. Clark, J. Ostell, K.D. Pruitt, and I. Karsch-Mizrachi. Genbank. Nucleic Acids Research, 48(D1):D84–86, 2020.

10. Uniprot Consortium. Uniprot: the universal protein knowledgebase in 2021. Nucleic Acids Research, 49(D1):D840–D489, 2020.

11. H.M Berman, J Westbrook, Z Feng, G Gilliland, T.N Bhat, H Weissig, and P. E. Bourne. The protein data bank. Nucleic Acids Research, 28(D1):D235–D242, 2000.

12. D.H. Parks, M. Chuvochina, C. Rinke, A.J. Mussig, P. Chaumeil, and P. Hugenholtz. Gtdb: an ongoing census of bacterial and archaeal diversity through a phylogenetically consistent, rank normalized and complete genome-based taxonomy. Nucleic Acids Research, 50(D1): D785–D794, 2021.

13. R. D. Hipp. Sqlite, 2020.

14. M. Bayer. Sqlalchemy, in The Architecture of Open Source Applications Volume II: Structure, Scale, and a Few More Fearless Hacks. Mountain View, Colorado, US, 2012.

15. P.J.A. Cock, T. Antao, J.T. Chang, B.A. Chapman, C.J. Cox, A. Dalke, I. Friedberg, T. Hamelryck, F. Kauff, B. Wilczynski, and M. J. L. de Hoon. Biopython: freely available python tools for computational molecular biology and bioinformatics. Bioinformatics, 25(11):1422–1423, 2009.

16. D.L. Wheeler, D.A. Benson, S. Bryant, K. Canese, D.M. Church, R. Edgar, S. Federhen, W. Helmberg, T.L. Madden, J.U. Pontius, G.D. Schuler, L.M. Schriml, E. Sequeira, T.O. Suzek, T.A. Tatusova, and L. Wagner. Database resources of the national centre for biotechnology information: Update. Nucleic Acids Research, 33(D1):D39–D45, 2005.

17. T. Cokelaer, D. Pultz, L. M. Harder, J. Serra-Musach, and J. Saez-Rodriguez. Bioservices: a common python package to access biological web services programmaticall. Bioinformat-ics, 19(24):3241–3242, 2013.

18. T. Hamelryck and B. Manderick. Pdb parser and structure class implemented in python. Bioinformatics, 19(17):2308–2310, 2003.

19. Federhen. The ncbi taxonomy database. Nucleic Acids Research, 40(D1):D136–D143, 2005.

20. B. Grüning, R. Dale, A. Sjödin, B. A. Chapman, J. Rowe, C. H. Tomkins-Tinch, R. Valieris, and J. Köster. Bioconda: sustainable and comprehensive software distribution for the life sciences. Nature Methods, 33(D1):D39–D45, 2018.

21. Python Software Foundation. Python package index - pypi, 2022.

22. V. Lombard, H. G. Ramulu, E. Drula, P. M. Coutinho, and B Henrissat. The carbohydrate-active enzymes database (cazy) in 2013. Nucleic Acids Research, 42(D1):D490–D495, 2014.

23. K. Barrett, C.J. Hunt, L. Lange, and A.S. Meyer. Conserved unique peptide patterns (cupp) online platform: peptide-based functional annotation of carbohydrate active enzymes. Nucleic Acids Research, 48(W1):W110–W115, 2020.

24. F.M. Razeq, E. Jurak, P.K. Stogios, R. Yan, M. Tenkanen, M.A. Kabel, W. Wang, and E.R. Master. A novel acetyl xylan esterase enabling complete deacetylation of substituted xylans. Biotechnology for Biofuels and Bioproducts, 11(74), 2018.

25. N. Konno, K. Igarashi, N. Habu, M. Samejima, and A. Isogai. Cloning of the trichoderma reesei cdna encoding a glucuronan lyase belonging to a novel polysaccharide lyase family. Applied and Environmental Microbiology, 75(1):101–107, 2009.

26. D. B. Singh and T. Tripathi. Frontiers in Protein Structure, Function, and Dynamics. Springer, Singapore, 1st edition, 2020.

27. E. Litte, P. Bork, and R. F. Doolittle. Tracing the spread of fibronectin type iii domains in bacterial glycohydrolases. Journal of Molecular Evolution, 39:631–643, 1994.

28. R. L. Szabady and R. A. Welch. Stce peptidase and the stce-like metalloendopeptidases. In N. D. Rawlings and G. Salvesen, editors, Handbook of Proteolytic Enzymes. Academic Press, Massachusetts, 3rd edition, 2013.

29. S. Adindla, K. K. Inampudi, K. Guruprasad, and L. Guruprasad. Identification and analysis of novel tandem repeats in the cell surface proteins of archeal and bacterial genomes using computational tools. Comparative and Functional Genomics, 5(1):2–16, 2004.

30. J. Hallgren, K. D. Tsirigos, M. D. Pedersen, J. J. A. Armenteros, P. Marcatili, H. Nielsen, A. Krogh, and O. Winther. Deeptmhmm predicts alpha and beta transmembrane proteins using deep neural network. BioRix, 2022.

31. M.L. Garron and B. Henrissat. The continuing expansion of cazymes and their families. Current Opinion in Chemical Biology, 53:82–87, 2019.

32. A. Ochiai, T. Itoh, Y. Maruyama, A. Kawamata, B. Mikami, W. Hashimoto, and K. Murata. Structural determinants responsible for substrate recognition and mode of action in family 11 polysaccharide lyases. The Journal of Biological Chemistry, 284(15):10181–10189, 2007.

33. A. Ochiai, T. Itoh, A. Maruyama, B. Mikami, W. Hashimoto, and K. Murata. Structural determinants responsible for substrate recognition and mode of action in family 11 polysaccharide lyases. The Journal of Biological Chemistry, 284(15):10181–10189, 2020.

34. R. Silva, C. Jers, H. Otten, C. Nyffenegger, D. M. Larsen, P. M. F. Derkx, A. S. Meyer, J. D. Mikkelsen, and S. Larsen. Design of thermostable rhamnogalacturonan lyase mutants from bacillus licheniformis by combination of targeted single point mutation. Applied Microbiology and Biotechnology, 98(10):4521–4531, 2014.

35. A. Mølgaard and S. Kauppienen, S. Larsen. Rhamnogalacturonan acetylesterase eluci-dates the structure and function of a new family of hydrolases. Structure, 8(4):373–383, 200.

36. A. Langkilde, S. M. Kristensen, L. L. Leggio, A. Mølgaard, J. H. Jensen, A. R. Houk, J. C. N. Poulsen, and S. Kauppinen, S. Larsen. Short strong hydrogen bonds in proteins: a case study of rhamnogalacturonan acetylesterase. Acta Crystallographic. Section D, Biological Crystallography, D64(8):851–863, 2008.

37. A. Levasseur, E. Drula, V. Lombard, P.M. Coutinho, and B. Henrissat. Expansion of the enzymatic repertoire of the cazy database to integrate auxiliary redox enzymes. Biotechnolgy Biofuels, 6(41), 2013.

38. K. Robak and M. Balcerek. Review of second generation bioethanol production from residual biomass. Food Technology and Biotechnology, 56(2):174–187, 2018.

39. D. Ndeh, A. Rogowski, A. Cartmell, A.S. Luis, A. Baslé, and J. et al Gray. Complex pectin metabolism by gut bacteria reveals novel catalytic functions. Nature, 544:65–70, 2017.

40. U. Avci, M. J. Pena, and M. A. O’Neil. Changes in the abundance of cell wall apiogalactur-onan and xylogalacturonan and conservation of rhamnogalacturonan ii structure during the diversification of the lemnoideae. Planta, 247:953–971, 2018.

41. I. Martínez-Martínez, J. Navarro-Fernández, J. D. Lozada-Ramírez, F. García-Carmona, and A. Sánchez-Ferrer. Yest: a new rhamnogalacturonan acetyl esterase from bacillus subtilis. Proteins, 71(1):379–388, 2008.

42. A. Ochiai, Y. Itoh, A. Kawamata, W. Hashimoto, and K. Murata. Plant cell wall degradation by saprophytic bacillus subtilis strains: Gene clusters responsible for rhamnogalacturonan depolymerization. Applied and Environmental Microbiology, 73(12):3803–3813, 2007.

43. S. Kauppinen, S. Christagu, L. V. Kofod, T. Halkier, K. D”orreich, and H. Dalbøge. Molecular cloning and characterization of a rhamnogalacturonan acetylesterase from aspergillus aculeatus: Synergism between rhamnogalacturonan degrading enzymes. Journal of Biological Chemistry, 270(45):27172–27178, 1995.

44. C. Montanier, A.L. van Bueren, C. Dumon, J.E. Flint, M.A. Correia, and J.A. Prates. Evidence that family 35 carbohydrate binding modules display conserved specificity but divergent function. Proceedings of the National Academy of Sciences, 106(9):3065–3070, 2009.

45. M. Mauri, T. Elli, G. Caviglia, G. Uboldi, and M. Azzi. Rawgraphs: A visualisation platform to create open outputs. In Proceedings of the 12th Biannual Conference on Italian SIGCHI Chapter, pages 1–5, 2017.

46. S. F. Altschul, W. Gish, W. Miller, E. W. Myers, and D. J. Lipman. Basic local alignment search tool. Journal of Molecular Biology, 215(3):403–410, 1990.

47. D. A. Rasko, G. S. A. Myers, and J. Ravel. Visualization of comparative genomic analyses by blast score ratio. BMC Bioinformatics, 6(2), 2005.

48. T. Galili, A. O’Callaghan, J. Sidi, and C. Sievert. heatmaply: an r package for creating interactive cluster heatmaps for online publishing. Bioinformatics, 34(9):1600–1602, 2018.

49. H. Wickham. ggplot2: Elegant Graphics for Data Analysis. Springer-Verlag, New York, 2016.

50. F. Teufel, J.J. Almagro Armenteros, A. R. Johansen, M. H. Gislason, S. I. Pihl, K. D. Tsirigos, O. Winther, S. Brunak, G. von Heijne, and H. Nielsen. Signalp 6.0 predicts all five types of signal peptides using protein language models. Nature Biotechnology, 40:1023–1025, 2022.

51. J. Jumper, R. Evans, and A. et al Pritzel. Highly accurate protein structure prediction with alphafold. Nature, 596:583–589, 2021.

52. E. F. Pettersen, T. D. Goddard, C. C. Huang, G. S. Couch, D. M. Greenblatt, E. C. Meng, and T. E. Ferrin. Ucsf chimera–a visualization system for exploratory research and analysis. Journal of Computational Chemistry, 25(13):1605–1612, 2004.

53. E. C. Meng, E. F. Pettersen, G. S. Couch, C. C. Huang, and T. E. Ferrin. Tools for integrated sequence-structure analysis with ucsf chimera. BMC Bioinformatics, 7(339), 2006.

54. M. Steinegger and J. Söding. Mmseqs2 enables sensitive protein sequence searching for the analysis of massive data sets. Nature Biotechnology, 35(11):1026–1028, 2017.

55. H. Zhange, T. Yohe, L. Huang, S. Entwistle, P. Wu, Z. Yang, P. K. Busk, Y. Xu, and Y. Yin. dbcan2: a meta server for automated carbohydrate-active enzyme annotation. Nucleic Acids Research, 46(W1):W95–W101, 2018.

56. J. Söding, A. Biegert, and A. N. Lupas. The hhpred interactive server for protein homology detection and structure prediction. Nucleic Acids Research, 33(Web Server Issue):W244–W248, 2005.

