## Additional Files for "cazy_webscraper: local compilation and interrogation of comprehensive CAZyme datasets": additional_file_6.pdf

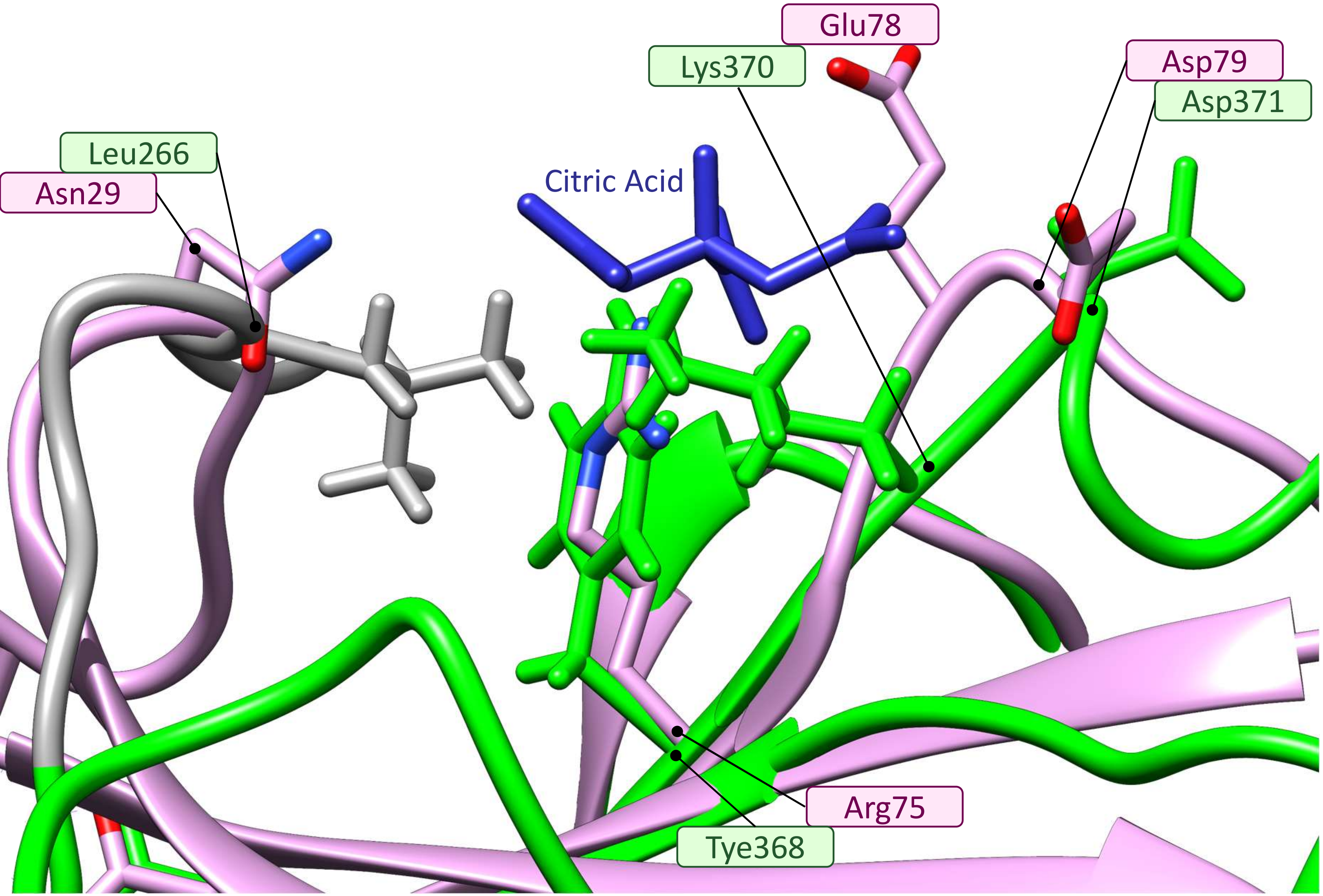


**Figure SI.1. Citric acid binding residues in PL20 enzyme PDB:2ZZJ and predicted structure fold of *HX109_05010*.**

*The structural fold of PDB:2ZZJ (shown in pink) superimposed onto the predicted structural fold (from Alphafold, v2.1.0) for the PL20 domain in HX109_05010 (shown in green, and disorganised region shown in grey). A citric acid in the PDB:2ZZJ structure is shown in blue. The side chains of residues in PDB:2ZZJ that bind citric acid and the corresponding residues in HX109_05010 are shown and annotated.*

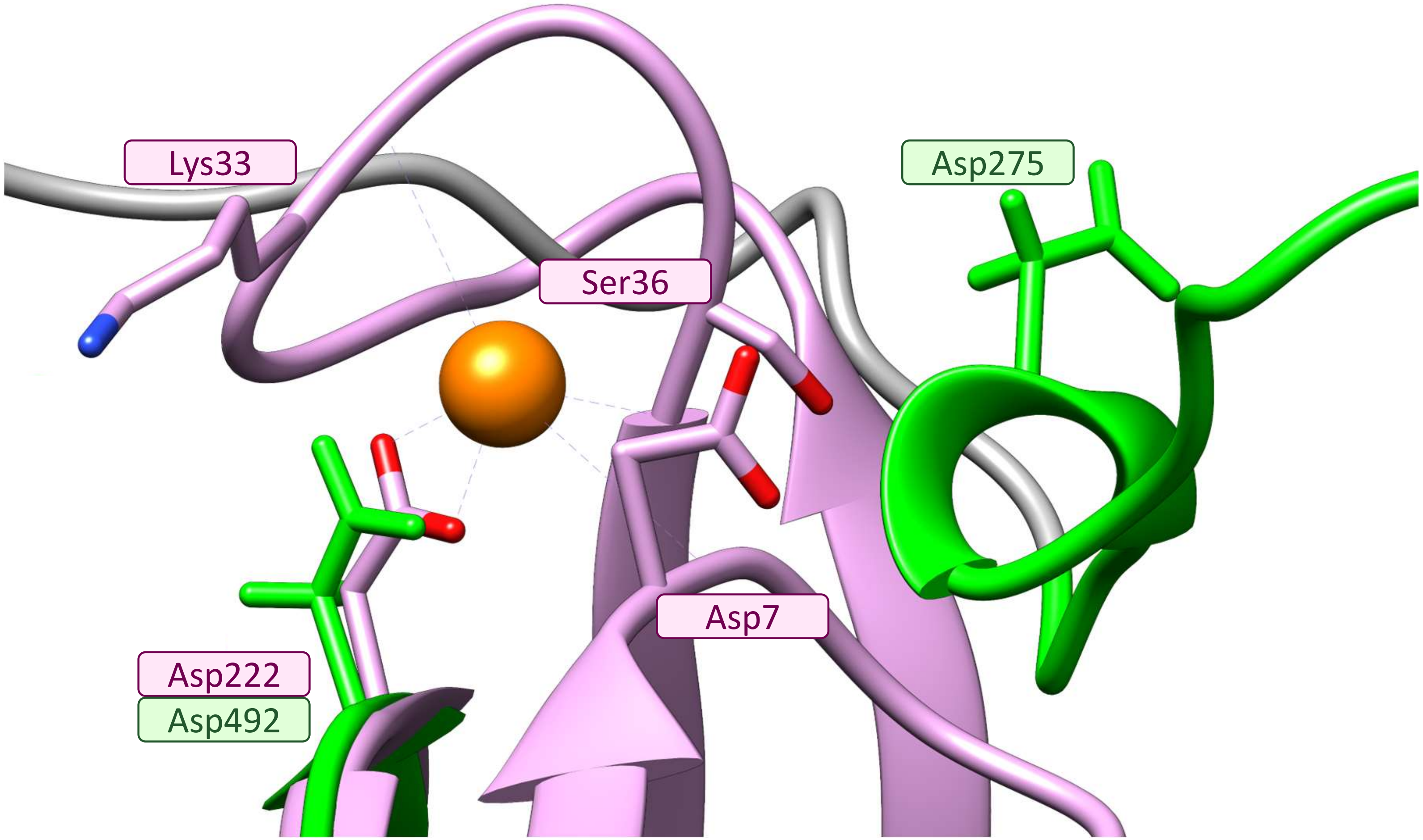


**Figure SI.2. Calcium ion binding residues in PL20 enzyme PDB:2ZZJ and predicted structure fold of *HX109_05010*.**

*The structural fold of PDB:2ZZJ (shown in pink) superimposed onto the predicted structural fold (from Alphafold, v2.1.0) for the PL20 domain in HX109_05010 (shown in green, and disorganised region shown in grey). A calcium ion in the PDB:2ZZJ structure is shown in orange. The side chains of residues in PDB:2ZZJ that bind calcium and the corresponding residues in HX109_05010 are shown and annotated.*

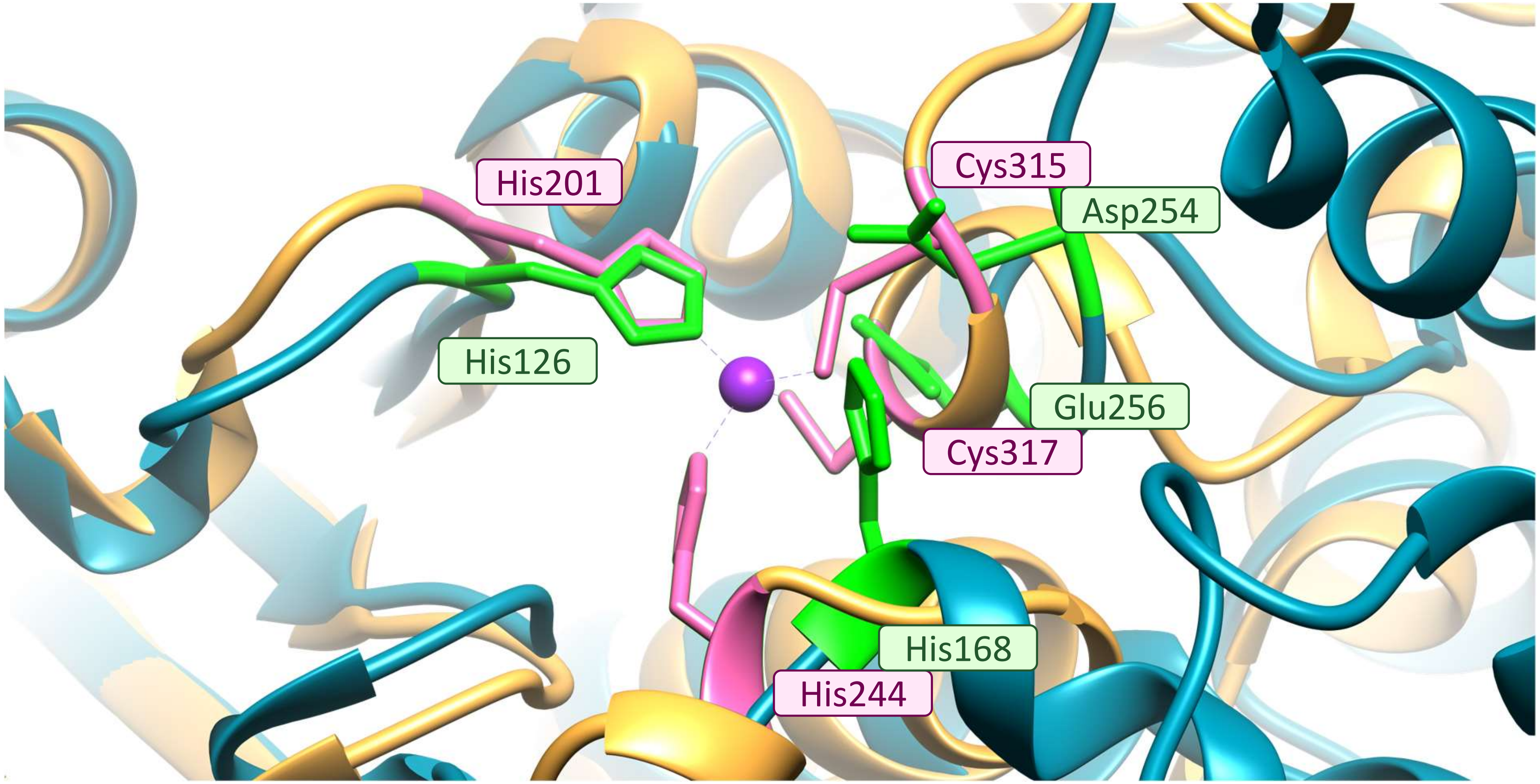


**Figure SI.3. Metal ion binding site in PDB:6GOC.** The metal ion (Zinc, shown in purple) binding site in PDB:6GOC (NCBI accession ALJ42174.1, shown in gold), superimposed onto the structural fold of *TBR22_41900* (NCBI accession BCS34995.1) CE19 domain (predicted by alphafold), shown in blue, using Chimera MatchMaker tool. 6GOC metal binding residues are shown in pink; corresponding residues in *TBR22_41900* are highlighted in green.

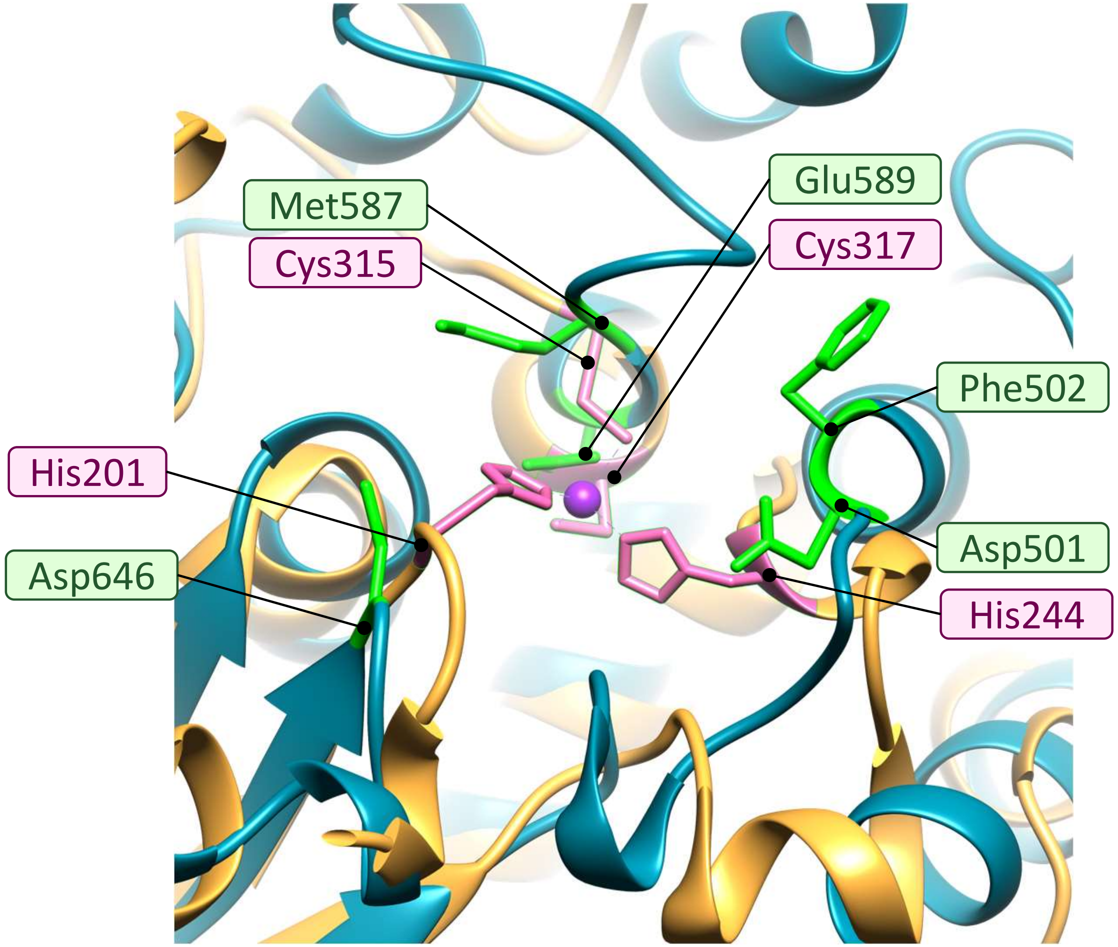


**Figure SI.4. Metal ion binding site in PDB:6GOC.** The metal ion (Zinc, shown in purple) binding site in PDB:6GOC (NCBI accession ALJ42174.1, shown in gold), superimposed onto the structural fold of *TBR22_41900* (NCBI accession BCS34995.1) second domain (predicted by alphafold), shown in blue, using Chimera MatchMaker tool.. 6GOC metal binding residues are shown in pink; corresponding residues in *TBR22_41900* are highlighted in green.

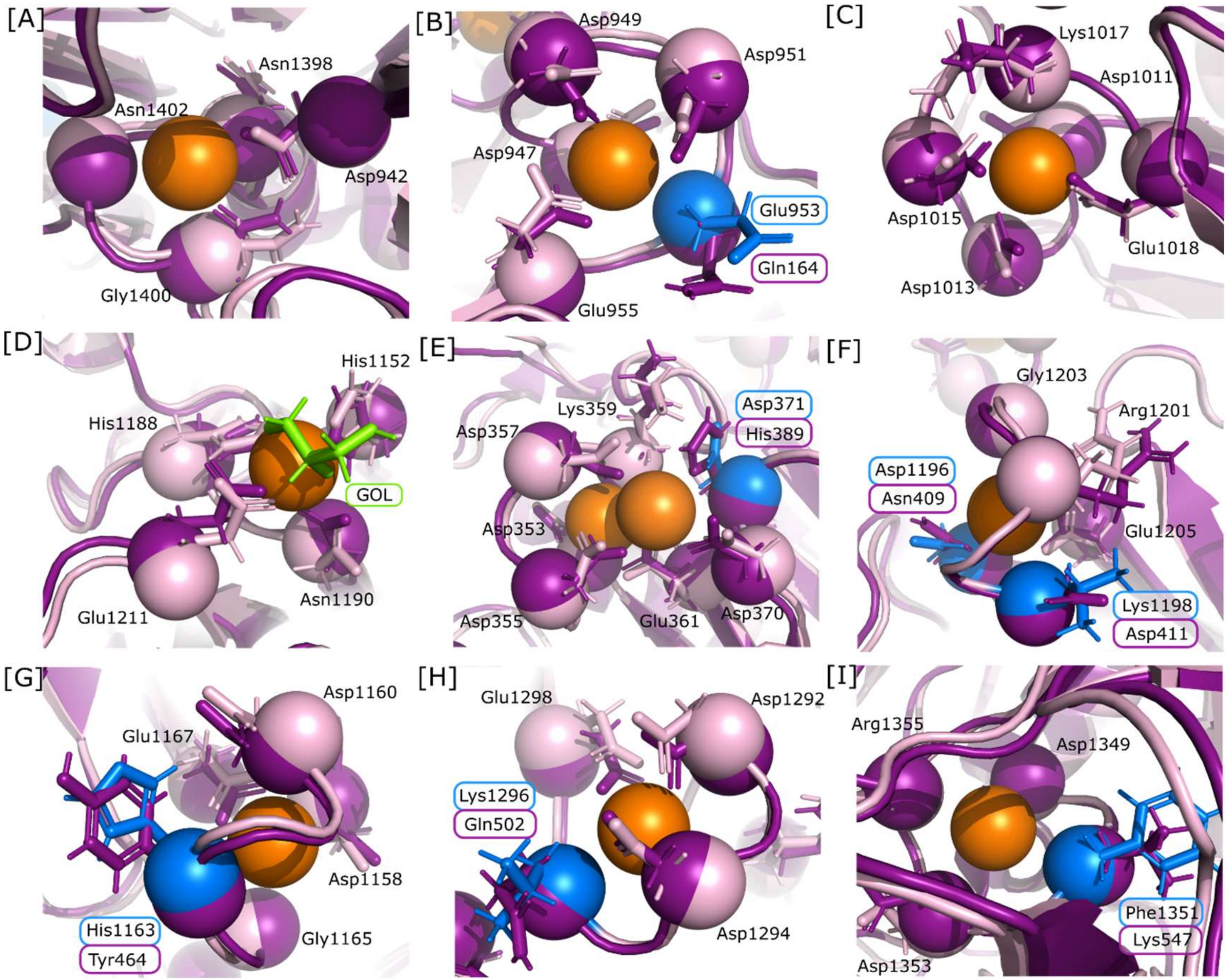


**Figure SI.5. Conserved calcium ion binding site residues in PL11 enzyme PDB:4CAG and predicted structural fold of *HUW50_16055*.**

*The structural fold of PDB:4CAG (shown in dark pink) superimposed onto the predicted structural fold (from Alphafold, v2.1.0) for the PL11 domain in HUW50_16055 (shown in light pink). Calcium ion binding residues in PDB:4CAG and corresponding residues in HUW50_16055 are shown as spheres with side chains shown as sticks. For residues that match, the position in HUW50_16055 is given. For residues that do not match, the residues in PDB:4CAG and HUW50_16055 are labelled, with the HUW50_16055 highlighted in blue.*

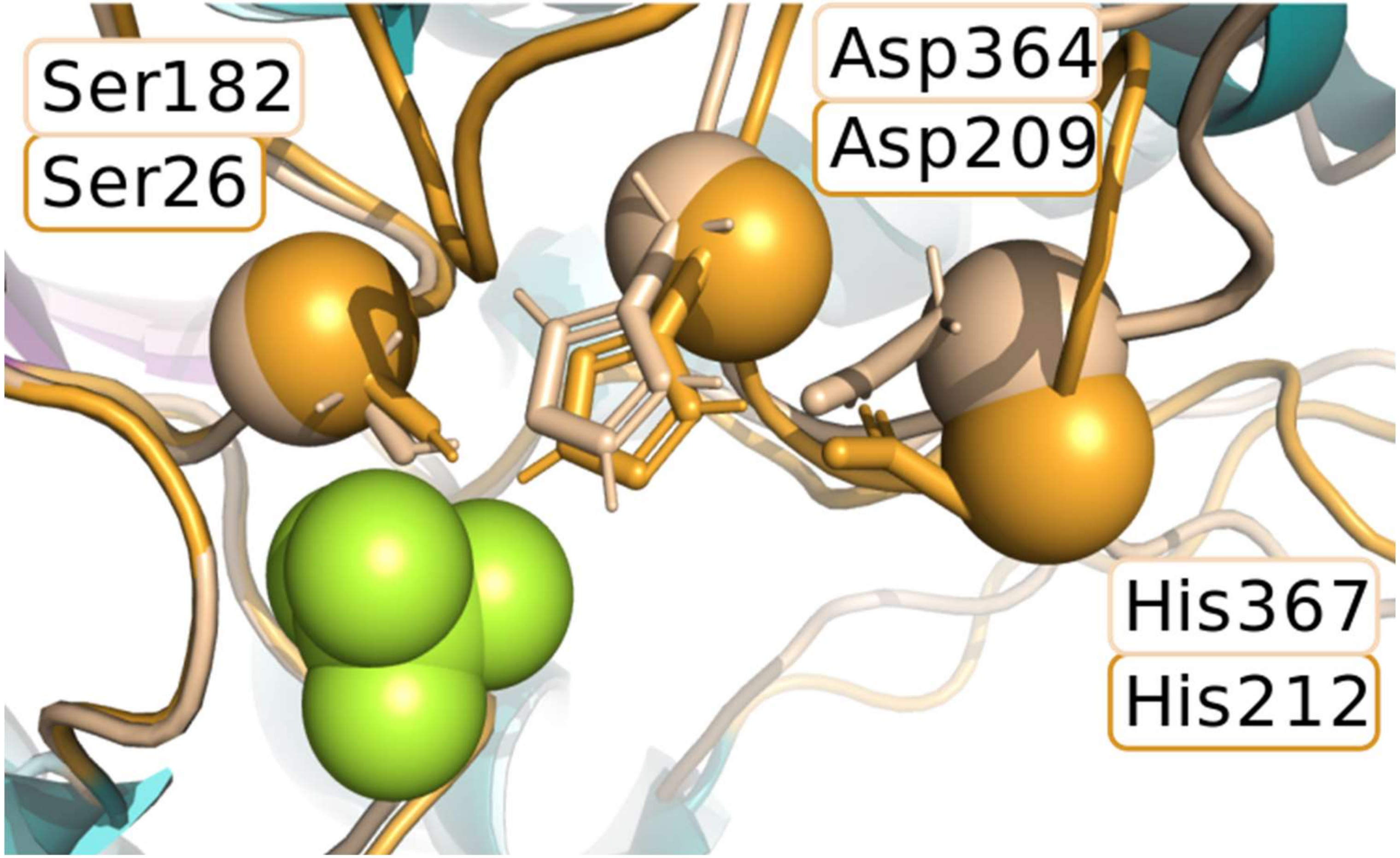


**Figure SI.6. Conserved binding site residues in CE12 enzyme PDB:1DEO and *HUW50_16055.***

*PDB:1DEO (shown in darker shades) superimposed onto the predicted structural fold (from Alphafold, v2.1.0) for the CE12 domai[illegible] in HUW50_16055 (shown in lighter shades). The catalytic triad in 1DEO and the corresponding residues in HUW50_16055 are shown as sphere with sidechains as sticks, and the residues are labelled.*
