## Additional Files for "cazy_webscraper: local compilation and interrogation of comprehensive CAZyme datasets": additional_file_17.docx

CE19 *TBR22_41900* Structure Comparison

Structural comparisons between the predicted fold of *TBR22_4900* (NCBI accession BCS34995.1) and hits from HHpred (accessed June 2022).

Structures obtained from RCSB-PDB were superimposed onto the predicted structures and Root Mean Square Distance (RMSD) calculated using the MatchMaker tool followed by the and Match-Align tool in UCSF Chimera (version 1.16) with default parameters. The superimpositions were visualised using PyMol (version 2.5.2).

### Sequence based superimposement of 6GOC onto TBR22

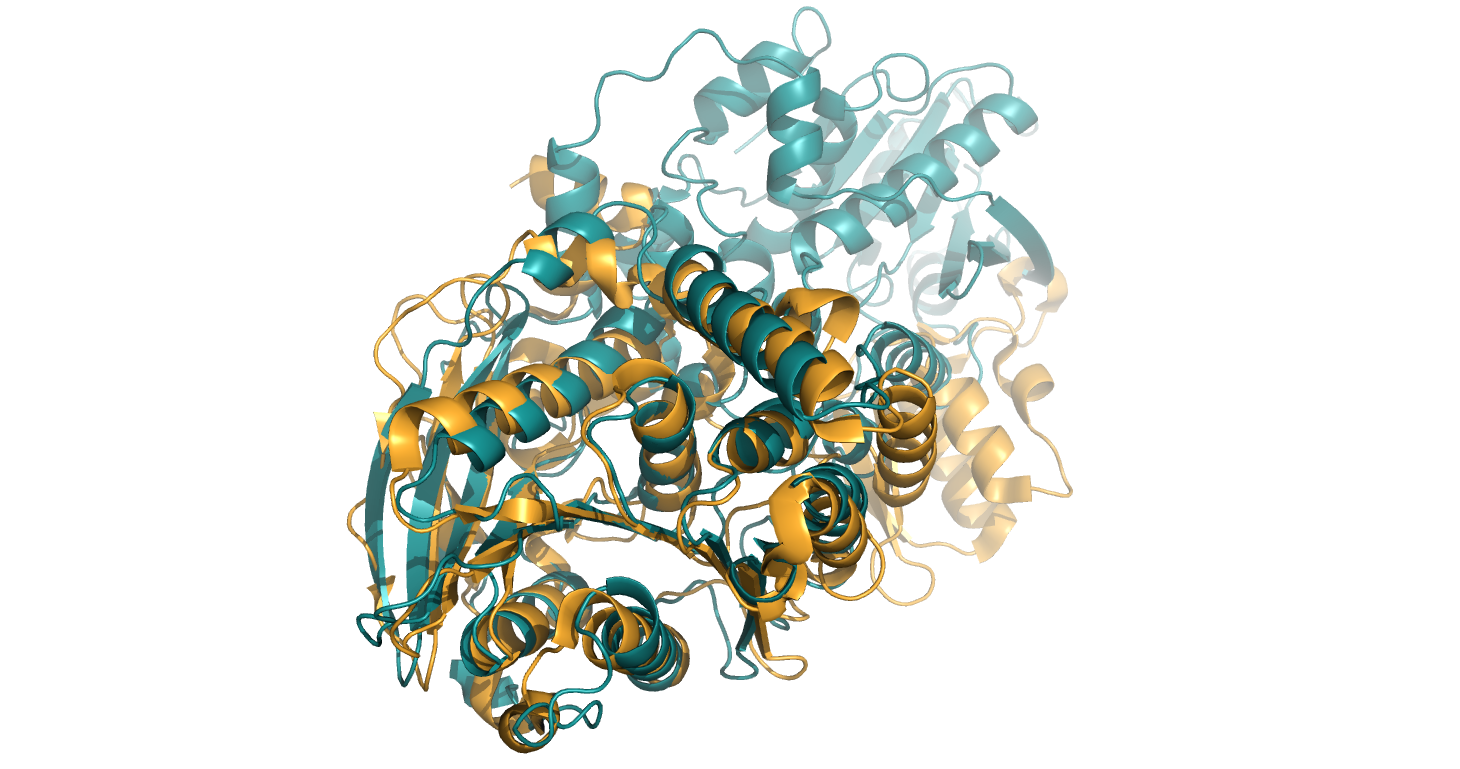

**TBR22 β-domain**

**CE19 domain**

The PDB 6GOC structural fold (shown in orange) superimposed onto the structural fold of TBR22 predicted by alphafold (version 2.1.0) shown as secondary structures in teal. RMSD: 2.048Å (across 311 alpha carbons). 6GOC was aligned onto the TBR22 CE19 domain.

### Sequence based superimposement of 3GY8 onto TBR22

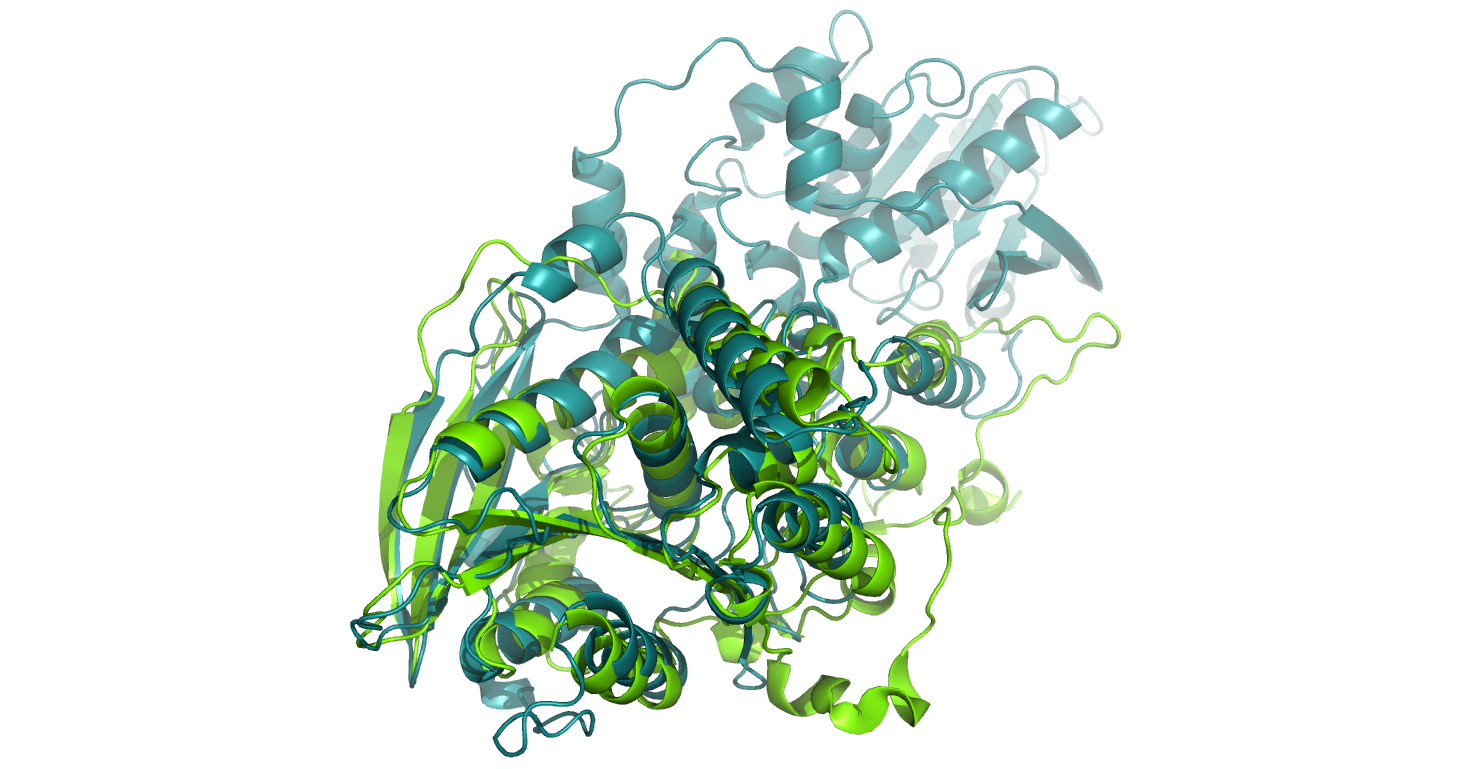

**CE19 domain**

**TBR22 β-domain**

The PDB 3GY8 structural fold (shown in green) superimposed onto the structural fold of TBR22 predicted by alphafold (version 2.1.0) shown as secondary structures in teal. RMSD: 1.950 Å (across 278 alpha carbons). 3GY8 was aligned onto the TBR22 CE19 domain.

### Sequence based superimposement of 3NUZ onto TBR22

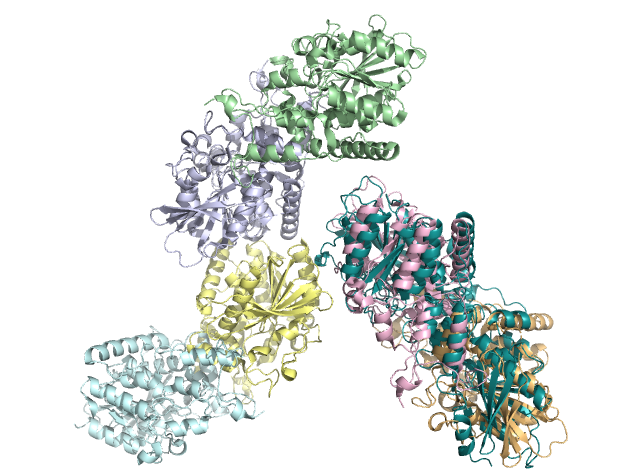

**TBR22**

The PDB 3NUZ structural fold (chain A shown in green, chain B in clue, chain C in yellow, chain D in blue, chain E in pink and chain F in orange) superimposed onto the structural fold of TBR22 predicted by alphafold (version 2.1.0) shown as secondary structures in teal. RMSD: 1.852 Å (across 275 alpha carbons).

**Command:** align 3nuz & chain E, selected_prediction

**Result:**

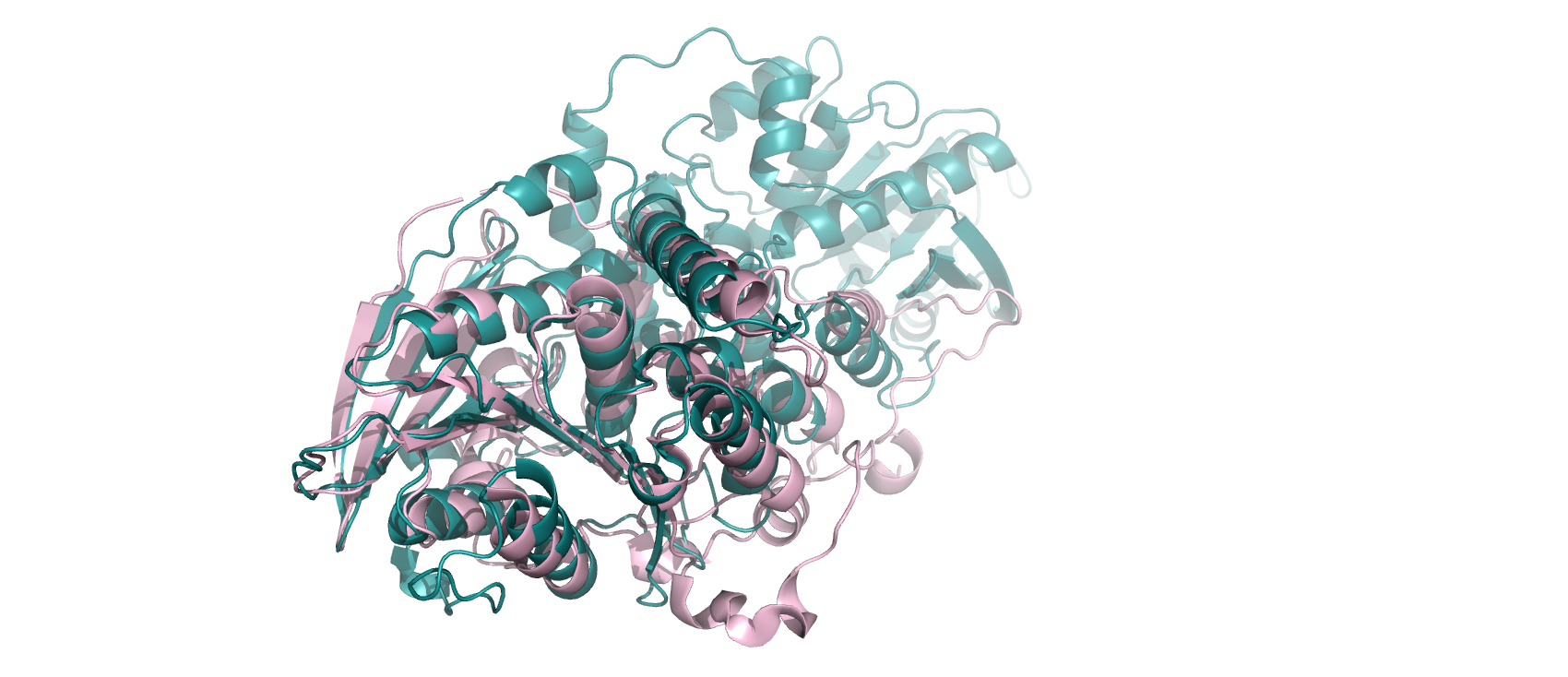

The PDB 3NUZ structural fold (chain E in pink) superimposed onto the structural fold of TBR22 predicted by alphafold (version 2.1.0) shown as secondary structures in teal. RMSD: 2.038 (863 to 863 atoms).

### Structure based superimposement of 6RUI onto TBR22

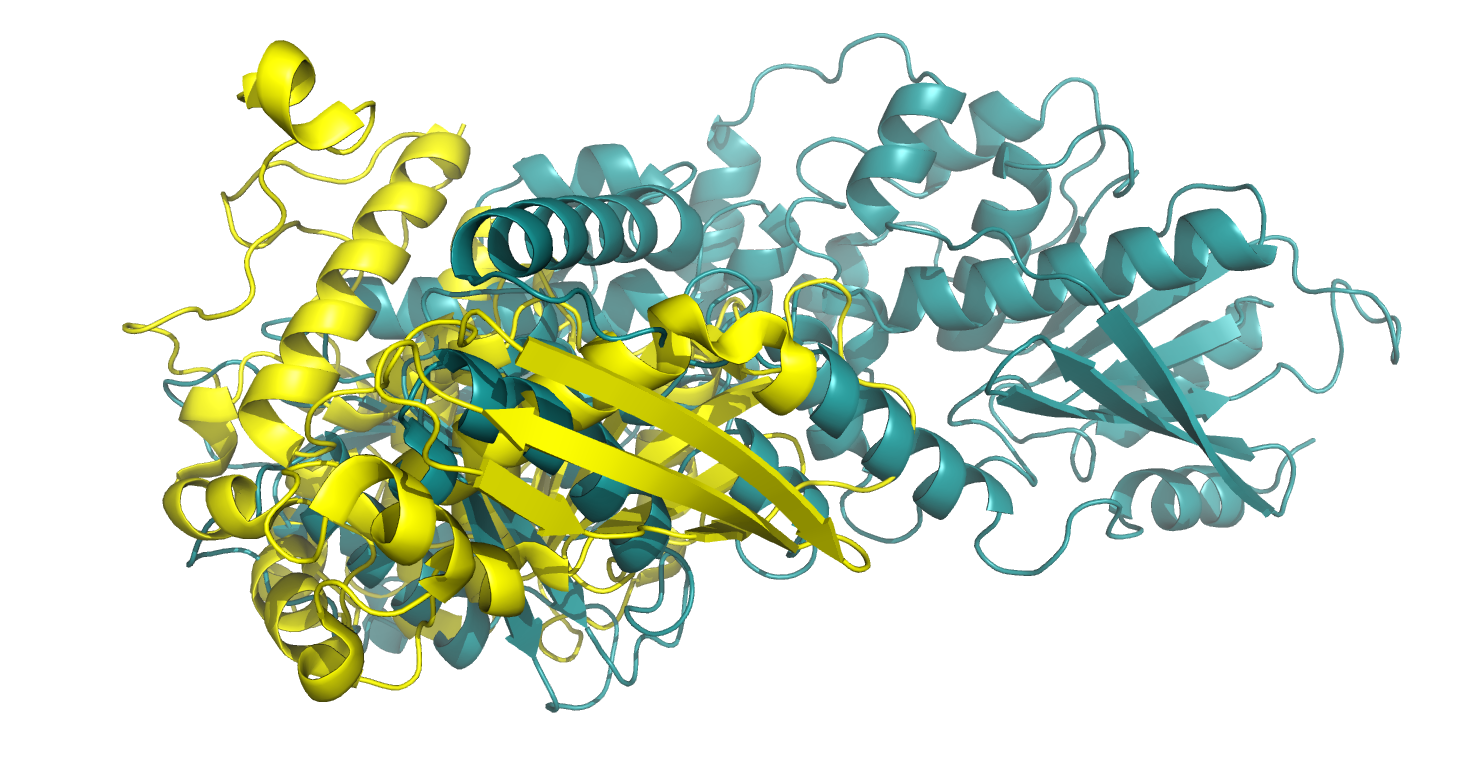

**CE19 domain**

**TBR22 β-domain**

The PDB 6RUI structural fold (shown in yellow) superimposed onto the structural fold of TBR22 predicted by alphafold (version 2.1.0) shown as secondary structures in teal. RMSD: 2.234 Å (across 231 alpha carbons).

### Sequence based superimposement of 6RUI onto TBR22 β-domain

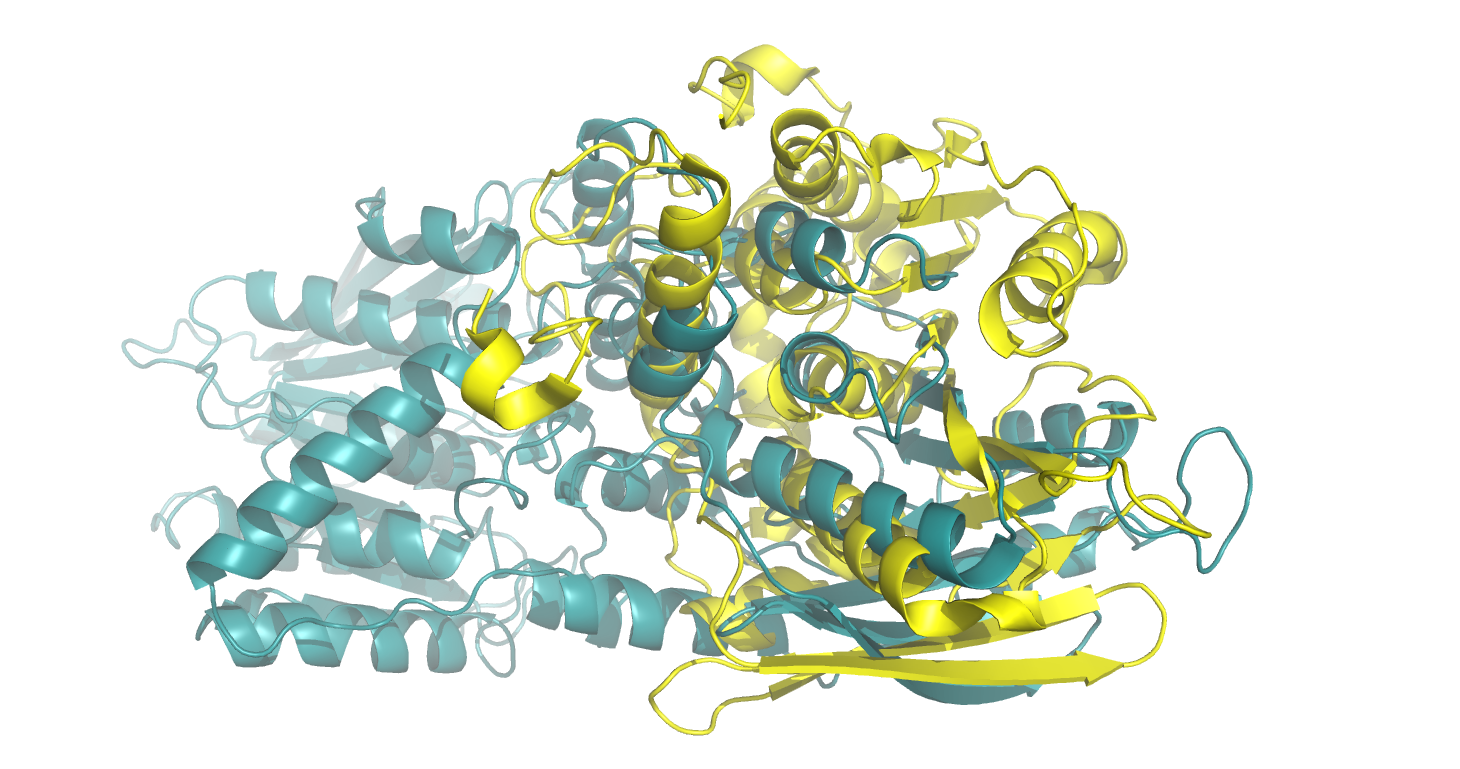

**CE19 domain**
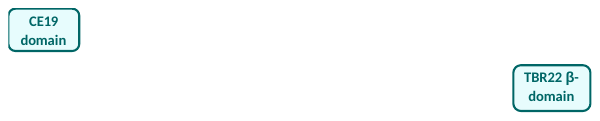

**TBR22 β-domain**

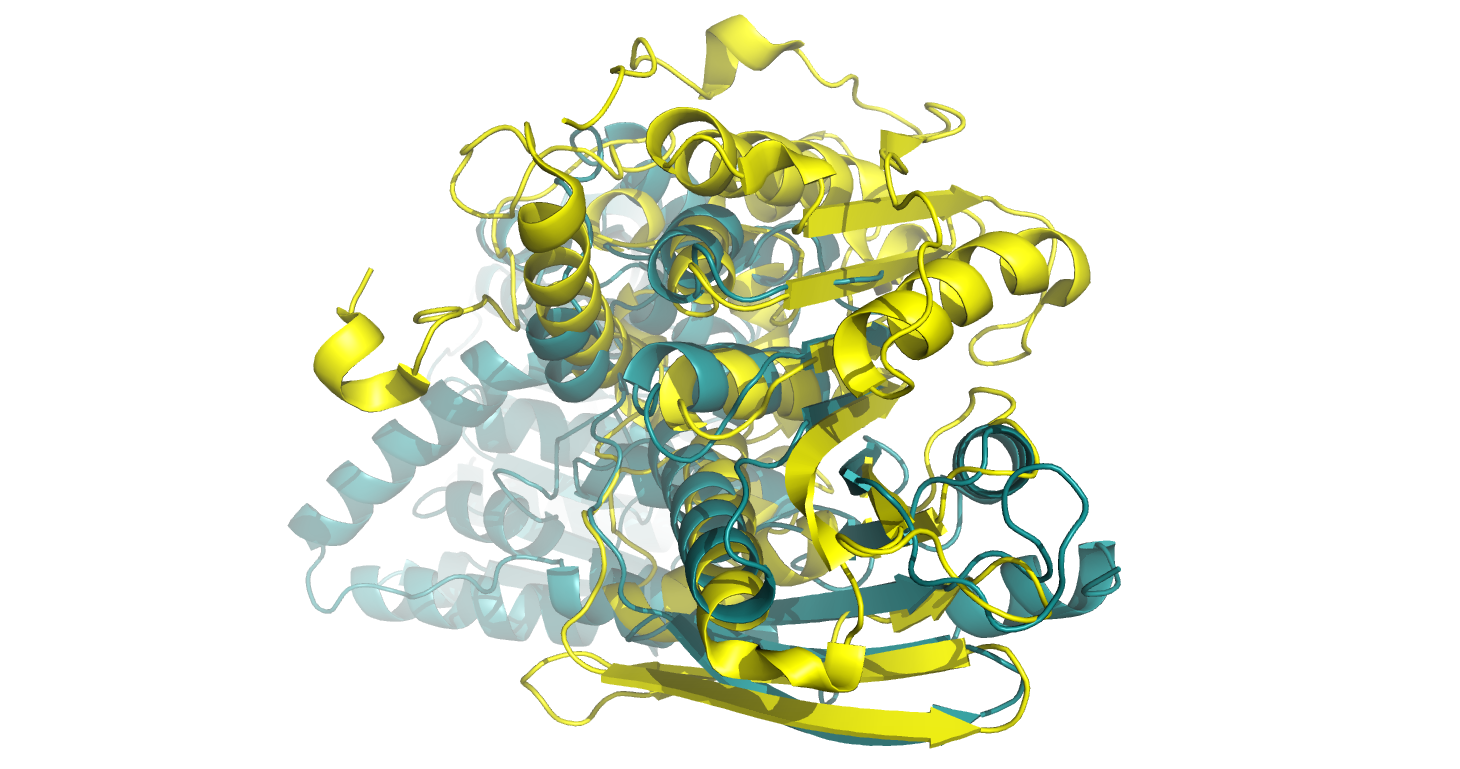

The PDB 6RUI structural fold (shown in yellow) superimposed onto the structural fold of TBR22 predicted by alphafold (version 2.1.0) shown as secondary structures in teal. RMSD: 2.352 Å (across 174 alpha carbons).
