## Supplementary figures and images for "cazy_webscraper: local compilation and interrogation of comprehensive CAZyme datasets"

### additional_file_2.pdf

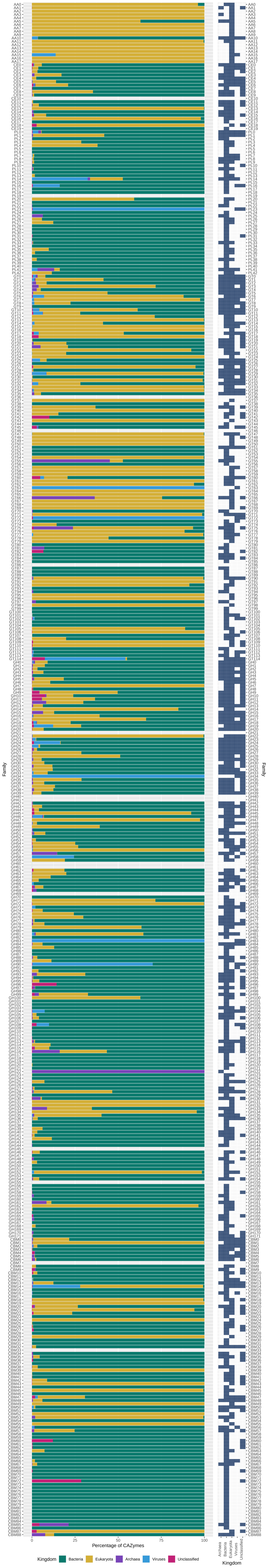
